# Within-host algal symbiont turnover is not a general response to environmental change in reef corals

**DOI:** 10.64898/2026.07.20.739650

**Authors:** Carly B. Scott

## Abstract

Flexibility in the coral-algal symbiosis is widely proposed as a mechanism of acclimatization to environmental change. Here, I compile colony-level longitudinal data spanning 67 coral species, 26 years, and 109 reefs globally to determine the predictors of within-host symbiont turnover. Across host species, environmental change showed no consistent association with turnover likelihood at either the symbiont genus or ITS2 sequencing variant level. Instead, resident symbiont genus was the strongest predictor of turnover, and transition probabilities between genera were often asymmetric. Host species differed in their turnover propensity under environmental change, with trait and phylogenetic similarity explaining most of this variation. Together, these results demonstrate that turnover is not necessarily a general response to environmental change across scleractinian coral species. Host traits, phylogeny, and symbiont identity may therefore help identify coral taxa most likely to undergo turnover as oceans warm, although its consequences for host acclimatization remain unresolved.

## Introduction

Symbiodiniaceae are a phylogenetically and functionally diverse family of dinoflagellate algae that form obligate endosymbioses with reef-building corals. At any given time, a single coral colony generally associates with just one dominant Symbiodiniaceae lineage, comprising > 99% of the *in hospite* community, while maintaining a diverse assemblage of rare background types at low abundance (Baker, 2003; Stat et al., 2009). However, which dominant symbiont a colony hosts is not fixed across its lifetime or uniform across a species. Conspecific colonies on the same reef can associate with functionally distinct symbiont genera (e.g., Innis et al., 2018), and symbiont community composition can vary across populations (e.g., Scott et al., 2025), depth gradients (e.g., Eckert et al., 2020), and time (e.g., Claar et al., 2020).

The flexibility of this relationship has been proposed as a mechanism enabling individual corals to acclimatize rapidly to changing environments (Buddemeier & Fautin, 1993), as Symbiodiniaceae exhibit broad functional and phylogenetic diversity (LaJeunesse et al., 2018). Since scleractinian corals can potentially associate with multiple genera or species, this flexibility may equip the host with a repertoire of partners advantageous under different environmental conditions. Under this view, elevated thermal stress or increased environmental variability should promote symbiont flexibility as corals acquire partners better suited to prevailing conditions. In response to perturbation, the dominant symbiont may be replaced by one of the background types (“shuffling”), or in rarer cases, by novel species acquired from the environment (“switching”; Baker, 2003; Baker et al., 2004; Boulotte et al., 2016).

Yet empirical evidence for symbiont turnover as a general stress response is mixed. Symbiont community composition is often remarkably stable across environmental gradients and stress events, even under severe thermal stress (Howells et al., 2020; LaJeunesse et al., 2010; Stat et al., 2009; Thornhill et al., 2009; Thornhill, Fitt, et al., 2006). Where turnover has been reported, patterns differ substantially across sampling sites and studies even within the same genus (e.g., *Siderastrea* spp.; Powell et al., 2026; Thornhill, Fitt, et al., 2006), suggesting that flexibility may be context-dependent rather than a general property of the symbiosis (Goulet, 2006).

Prior studies have found that coral-symbiont flexibility correlates with host phylogeny and geographic range (Swain et al., 2021; Zarate et al., 2024). However, these studies characterize flexibility at the reef or species scale, and propensity of an individual coral colony to experience symbiont community turnover remains unclear. Studies tracking symbiont community composition at the reef level often cannot fully disentangle population-scale processes, such as selective mortality of colonies hosting less thermally tolerant symbionts, from true within-colony turnover (e.g., Quigley et al., 2022; Van Nynatten et al., 2025).

Within-host symbiont dynamics may be additionally shaped by competitive interactions among Symbiodiniaceae strains. During early coral life stages, priority effects and other ecological dynamics can influence which Symbiodiniaceae lineage achieves dominance (McIlroy et al., 2020). More broadly, observed shifts in community composition during perturbation may reflect competitive release of previously suppressed background types as much as host-mediated selection for more advantageous partners (Abbott et al., 2021; Pettay et al., 2015; Scott et al., 2024). Thus, variation in turnover propensity across species may reflect asymmetries in competitive dynamics among resident symbiont lineages in addition to differences in host biology.

Here, I compile longitudinal data spanning 67 coral species, 26 years, and 109 globally distributed reefs to test the hypothesis that variation in symbiont turnover propensity in response to changing environments is not universal but is instead structured by host phylogeny, morphological traits, and symbiont-symbiont interactions.

## Materials & Methods

### Identification of studies for inclusion

Prior to searching, I defined three criteria for study inclusion. Studies were required to have measured the dominant symbiont of the same coral colony at least twice through time, generating colony-level, longitudinal time series. Measurements must have been conducted in situ on the reef rather than in aquaria or laboratory settings. Finally, I excluded colonies that underwent experimental manipulation, including reciprocal transplants and common garden treatments. This final criteria was applied to focus on turnover as colonies experienced environmental change on their source reef, as change in exposure between samples from reciprocal transplants can reflect both spatial differences between reefs and environmental change through time complicating the modeling approach.

Searches were conducted from May to June 2026 across two databases, PubMed and Google Scholar. PubMed was searched using a targeted Boolean query string for longitudinal symbiont dynamics in Scleractinia (Table S1), returning 201 results. Google Scholar was additionally searched using two broader query strings (Table S1), returning 591 additional results. Google Scholar results were screened manually and references cited within retained papers were mined to identify further eligible studies. In October 2026, a supplemental search was conducted using ChatGPT Sol5.6 model to identify additional manuscripts which may have been missed according to the search criteria (full chat log in Supplemental File 3). Additionally, data listed online in the SymPortal.org database were screened but did not yield any additional published manuscripts not already identified by other methods. After applying the inclusion criteria described above, 42 studies were retained (Table 1), collectively spanning 2,996 colonies from 67 scleractinian coral species sampled between 1998 and 2024 across 109 globally distributed reefs (Figure 1).

**Figure 1.**
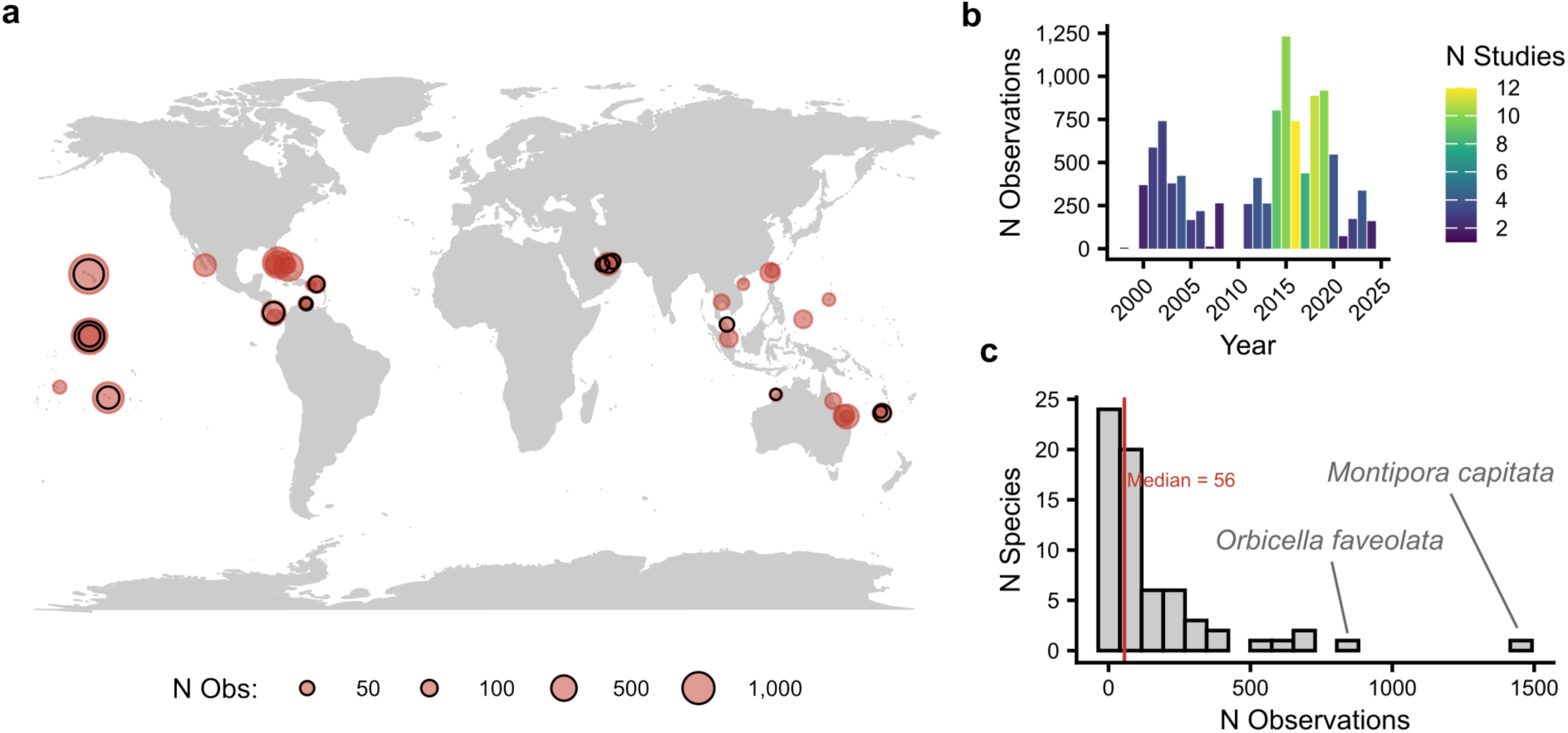
Overview of included data. (a) Geographic scope of studies included in this meta-analysis. Studies spanned the Indo-Pacific, Caribbean/Western Atlantic, Red Sea, and Eastern Tropical Pacific. Black outlines indicate locations included in the DIV-level analysis (b) Number of symbiont community observations by sampling year. Bar color indicates how many studies observations were sourced from. (c) Histogram of the number of observations for each coral species included in this study. Species had a median of 56 observations. The most represented species, *Montipora capitata* and *Orbicella faveolata,* are indicated.

**Table 1.** Summary of studies included in meta-analysis, indicating the total number of coral species (SPECIES), individual colonies (COLONIES), total observations through time (OBS), region and number of sites observed (REGION/SITES), and temporal extent of study (DATES).

| STUDY | SPECIES | COLONIES | OBS | REGION (SITES) | DATES | DIV |
|---|---|---|---|---|---|---|
| **(CHEN ET AL., 2005)** | 1 | 6 | 42 | NE Asia (1) | 2000-2001 |  |
| **(THORNHILL, FITT, ET AL., 2006)** | 6 | 126 | 1728 | Northern Caribbean, Florida & Bahamas (3) | 1998-2004 |  |
| **(THORNHILL, LAJEUNESSE, ET AL., 2006)** | 3 | 24 | 150 | Northern Caribbean, Florida & Bahamas (7) | 2002-2005 |  |
| **(JONES ET AL., 2008)** | 1 | 127 | 254 | Great Barrier Reef (1) | 2004-2006 |  |
| **(SAMPAYO ET AL., 2008)** | 1 | 7 | 20 | Great Barrier Reef (1) | 2005-2006 |  |
| **(STAT ET AL., 2009)** | 10 | 139 | 417 | Great Barrier Reef (1) | 2001-2002 |  |
| **(THORNHILL ET AL., 2009)** | 2 | 50 | 200 | Northern Caribbean, Florida & Bahamas (5) | 2003-2005 |  |
| **(LAJEUNESSE ET AL., 2010)** | 1 | 121 | 319 | Eastern Tropical Pacific (2) | 2006-2008 |  |
| **(CHEN ET AL., 2011)** | 1 | 3 | 21 | NE Asia (1) | 2008 |  |
| **(KIMES ET AL., 2013)** | 1 | 6 | 12 | SE Caribbean (1) | 2007 |  |
| **(HUME ET AL., 2015)** | 6 | 18 | 144 | NW Arabian Sea (1) | 2011-2013 |  |
| **(ROUZÉ ET AL., 2016)** | 1 | 11 | 84 | Central South Pacific (2) | 2011-2012 |  |
| **(CUNNING ET AL., 2016)** | 1 | 60 | 337 | Hawaiian Islands (3) | 2014-2015 |  |
| **(MANZELLO ET AL., 2019)** | 1 | 124 | 248 | Northern Caribbean, Florida & Bahamas (3) | 2015-2016 |  |
| **(CARBALLO-BOLAÑOS ET AL., 2019)** | 1 | 25 | 75 | NE Asia (2) | 2016-2017 |  |
| **(EPSTEIN ET AL., 2019)** | 1 | 10 | 88 | Great Barrier Reef (1) | 2016-2017 |  |
| **(ROUZÉ ET AL., 2019)** | 3 | 45 | 360 | Central South Pacific (4) | 2011-2012 |  |
| **(THOMAS ET AL., 2019)** | 2 | 9 | 36 | SW Pacific (4) | 2015-2016 |  |
| **(CLAAR ET AL., 2020)** | 1 | 76 | 238 | Eastern Micronesia (14) | 2014-2019 | Y |
| **(HOWELLS ET AL., 2020)** | 3 | 118 | 375 | NW Arabian Sea (4) | 2012-2014 | Y |
| **(JUNG ET AL., 2021)** | 1 | 15 | 30 | Timor Sea (1) | 2016 | Y |
| **(JANDANG ET AL., 2022)** | 2 | 32 | 81 | SE Asia (1) | 2018 |  |
| **(LEINBACH ET AL., 2023)** | 1 | 26 | 50 | Central South Pacific (1) | 2019 | Y |
| **(MCRAE ET AL., 2023)** | 3 | 30 | 111 | NE Asia (2) | 2018-2019 |  |
| **(NÚÑEZ-PONS ET AL., 2023)** | 3 | 127 | 501 | Hawaiian Islands (1) | 2014-2015 | Y |
| **(PALACIO-CASTRO ET AL., 2023)** | 3 | 29 | 87 | Eastern Tropical Pacific (1) | 2014-2016 |  |
| **(QUEK ET AL., 2023)** | 3 | 24 | 144 | SE Asia (2) | 2018-2020 |  |
| **(STARKO ET AL., 2023)** | 1 | 163 | 524 | Eastern Micronesia (14) | 2014-2019 | Y |
| **(DE SOUZA ET AL., 2023)** | 1 | 468 | 936 | Hawaiian Islands (1) | 2018-2019 | Y |
| **(CUNNING ET AL., 2024)** | 9 | 56 | 141 | SE Caribbean (3) | 2017-2019 | Y |
| **(LOCK ET AL., 2024)** | 1 | 19 | 30 | Western Micronesia (1) | 2018 |  |
| **(TERRANEO ET AL., 2024)** | 4 | 62 | 180 | SW Pacific (4) | 2016 | Y |
| **(ALESSI ET AL., 2024)** | 1 | 21 | 52 | SW Pacific (2) | 2021 | Y |
| **(LI ET AL., 2025)** | 1 | 5 | 20 | SE Asia (1) | 2023-2024 | Y |
| **(TURNHAM ET AL., 2025)** | 13 | 32 | 146 | Western Micronesia (1) | 2013-2022 |  |
| **(VAN NYNATTEN ET AL., 2025)** | 2 | 237 | 605 | Eastern Micronesia (16) | 2013-2023 | Y |
| **(POWELL ET AL., 2026)** | 3 | 33 | 81 | SE Caribbean (4) | 2020-2021 | Y |
| **(GLASS ET AL., 2026)** | 1 | 163 | 440 | Mesoamerica (4) | 2022-2024 | Y |
| **(BUZZONI ET AL., 2026)** | 1 | 237 | 599 | Eastern Micronesia (12) | 2014-2023 | Y |
| **(LEE ET AL., 2026)** | 1 | 10 | 56 | SE Asia (1) | 2019-2020 | Y |
| **(HUFFMYER ET AL., 2026)** | 3 | 125 | 421 | Central South Pacific (3) | 2020 | Y |
| **(SCOTT ET AL., 2026)** | 1 | 7 | 25 | SE Caribbean (2) | 2022 |  |
| **TOTAL** | **67** | **2996** | **10398** | **15 Regions, 109 Sites** | **1998-2024** |  |

### Genus-level data extraction & cleaning

For each study, I extracted the dominant symbiont genus directly from supplemental relative abundance data if available. A symbiont genus was classified as dominant if it comprised greater than 90% of the *in hospite* community; if two or more genera each exceeded 10% relative abundance, the colony was classified as co-dominant. This threshold was chosen to parallel the approximate detection limit of DGGE-based methods utilized in many early studies (~10%; Boulotte et al., 2016; Thornhill, LaJeunesse, et al., 2006), such that co-dominance classifications are conservative and less likely to reflect methodological noise. For studies where data were available only as figures, dominant symbiont proportions were estimated from stacked bar charts using the 10% threshold assessed by bar length.

### DIV-level data extraction & cleaning

To look at finer-scale differences in symbiont turnover, a Defining Intragenomic Variant (DIV) level analysis was conducted for studies that used the SymPortal framework (Hume et al., 2019; Table 1; Figure 1a). This dataset included 16 studies and 3,127 observations from 1,766 colonies spanning 2012-2024 from 30 coral species. When available, the authors’ SymPortal DIV relative abundance table was used for the subsequent analysis. However, many studies only had raw reads accessible. For these studies, SymPortal was run locally on the University of North Carolina Longleaf computing cluster on reads downloaded from BioProject accessions. Additional details on which studies were processed in this way and the full bioinformatic pipeline can be found in script ‘00a_RawReads_SymPortal_Preprocessing.sh’ in the accompanying GitHub and Zenodo repositories.

For each coral with DIV-level data, I calculated the Bray–Curtis distance between symbiont communities across timepoints from the relative abundance of each DIV. UniFrac distances were also calculated for a sensitivity analysis (Supplemental File 2). Full details of data cleaning and dominant symbiont classification are available in scripts 00a-00c in the associated GitHub and Zenodo repositories. The final curated database can additionally be found in these repositories (https://github.com/cb-scott/longitudinal_symbiodiniaceae and Zenodo upon acceptance).

### Environmental data acquisition and dimensionality reduction

Environmental data were extracted for each unique combination of sampling location and date in the dataset. Sea surface temperature (SST), degree heating weeks (DHW), and SST anomaly were obtained from the NOAA Coral Reef Watch CoralTemp dataset (NOAA_DHW) via the ‘rerddap’ package in R. Ocean color variables, including chlorophyll-a concentration and diffuse attenuation coefficient (Kd490, a proxy for turbidity), were obtained from the ESA Ocean Color Climate Change Initiative version 6 dataset (occci_V6_8day_4km). Precipitation data were obtained from the IMERG monthly global precipitation dataset. For each sampling point, environmental variables were extracted over the calendar month of sampling when exact dates were unavailable, or over the 30 days preceding and including the sampling date when exact dates were known. Environmental data were retrieved within 1° of each sampling location, and the three nearest grid cells containing valid observations were selected. Despite the broad initial search window, the maximum distance between a sampling location and its selected environmental grid cells was 12 km. These spatially averaged observations were then summarized across the extraction period to calculate temporal means, minima, maxima, and standard deviations, as well as cumulative SST anomalies.

To reduce the dimensionality of the environmental data and address collinearity among variables, a principal components analysis (PCA) was performed on the full set of derived environmental metrics after standardization to zero mean and unit variance. The first three principal components (PC1, PC2, PC3) were retained for use as covariates in all subsequent models, collectively explaining 71.5% of environmental variance. Higher PC1 scores represented warmer, somewhat wetter, less turbid, and less variable conditions; higher PC2 scores represented anomalously warm, wetter, more turbid, and generally more variable conditions; and higher PC3 scores represented higher minimum temperatures, lower thermal variability, and weaker SST anomalies (Figure S1).

### Genus-level Bayesian hierarchical modeling

The primary goal of this analysis was to determine: (1) whether changes in environmental predictors increased the probability and extent of symbiont turnover generally, and (2) whether host phylogeny or morphological traits explained species-level variation in turnover propensity. These joint goals motivated the development of two complementary models, one at the genus-level and one at the DIV-level.

Using the genus-level dataset encompassing all 42 studies, I constructed a cumulative ordinal Bayesian hierarchical model to evaluate symbiont turnover. Turnover was classified into three ordered categories reflecting increasing departure from baseline symbiosis: no turnover (0), emergence of co-dominance between genera (1), and complete genus-level replacement (2). Dominance was defined as a single genus exceeding 90% relative abundance, while co-dominance occurred when two or more genera each exceeded 10% relative abundance. These thresholds were also chosen to distinguish meaningful changes in community composition from background diversity dynamics (e.g., Abbott et al., 2021) and reflect DGGE-based detection thresholds.

The model included fixed effects for log-transformed sampling interval duration (Δ*t*), the change in environmental principal components between subsequent sampling times for each colony (Δ*PC*1, Δ*PC*2, Δ*PC*3), and the prior dominant symbiont genus of each colony (*Prior Genus*), with *Cladocopium* as the reference level. To partition host species-level variance in baseline turnover propensity and environmental sensitivity across phylogenetic, trait-based, and residual components, the model included three sets of species-level random effects, each with random slopes for environmental Δ*PC*1, Δ*PC*2, and Δ*PC*3. First, a phylogenetic random effect was structured by a covariance matrix derived from a maximum clade credibility tree (Gault et al., 2021), capturing variance attributable to shared evolutionary history. Second, a trait-distance random effect was structured by a covariance matrix derived from Euclidean distances in trait space derived from McWilliam et al. (2018), capturing variance attributable to trait similarity among species. Third, a residual species-level random effect captured idiosyncratic species variation not explained by phylogeny or traits. Nested random intercepts for study, sampling site, and colony were included to account for non-independence of repeated measurements within colonies at sites and methodological differences across studies. An overview of the model structure is given below:

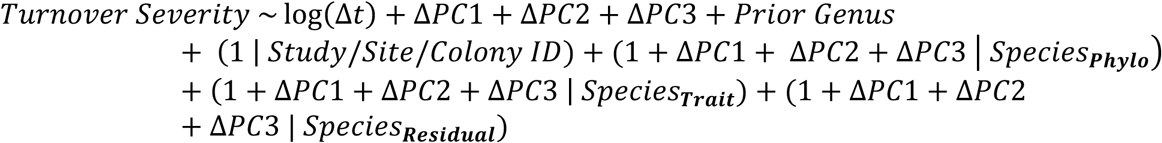

The phylogenetic covariance matrix was computed from a pruned and ultrametrized maximum clade credibility tree using ape::vcv.phylo with correlation scaling. Three species absent from the source tree (*Pocillopora acuta*, *Acropora downingi*, and *Pocillopora capitata*) were grafted manually as sister taxa to their closest relatives prior to pruning (respectively: *Pocillopora damicornis, Acropora aspera*, and *Pocillopora elegans*).

The trait covariance matrix was constructed as T = 1 − (D / max(D)), where D is the Euclidean distance matrix computed across trait principal components, and was projected to the nearest positive-definite matrix using Matrix::nearPD. An annotated biplot of coral trait PCs from McWilliam et al. (2018) is given in Figure S2.

Priors were specified as follows. Ordinal thresholds received normal priors centered at empirical cumulative log-odds values (1.81, 3.13) each with a standard deviation of 1.0, reflecting the strongly right-skewed distribution of turnover events. Fixed effect coefficients received normal(0, 1) priors. Total species-level variance parameters received lognormal(log(1), 1) priors, while the proportional contributions of phylogenetic, trait-associated, and residual species effects received Dirichlet(1, 1, 1) priors. Study-, site-, and colony-level standard deviations received half-Student-t(3, 0, 2.5) priors, and correlation matrices received LKJ(1) priors. Prior predictive checks confirmed that these priors produced plausible turnover rate distributions prior to model fit.

A key goal of the Bayesian analysis was to determine what proportion of species-level variation in turnover propensity is attributable to shared evolutionary history (phylogeny), shared traits, or idiosyncratic species differences not captured by either. To estimate these proportions directly rather than deriving them post-hoc from model parameters, variance partitioning was implemented in a custom Stan model via cmdstanr. Rather than fitting independent standard deviations for each random effect component and computing their relative contributions after the fact, this model parameterized the partition explicitly: for each model coefficient, a total species-level variance was sampled alongside a simplex vector that allocated that variance across the three components. The symmetric Dirichlet(1, 1, 1) prior is uniform over possible variance allocations, allowing any component to dominate without favoring one a priori. This parameterization allowed the relative contributions of phylogeny, traits, and residual species differences to be estimated jointly, with uncertainty quantified directly from their posterior distributions.

The model was fit with 4 chains, 2000 iterations (1000 warmup), adapt_delta = 0.95, and seed = 42. Convergence was assessed via R-hat statistics (R-hat < 1.01) and effective tail sample sizes (ESS > 400).

### DIV-level Bayesian hierarchical modeling

To evaluate finer-scale symbiont community turnover, I fit a complementary Bayesian hierarchical model using Bray–Curtis distances between DIV-level communities as a continuous response. Because Bray–Curtis distances are bounded between 0 and 1, I used a beta likelihood with a logit link. Distances equal to 1 (this only occurred for 2 observations) were adjusted to fall within the open unit interval by replacing them with the midpoint between 1 and the largest observed distance below 1.

The model otherwise followed the same hierarchical structure as the genus-level ordinal model, including fixed effects for sampling interval duration and changes in environmental conditions, nested study, site, and colony random intercepts, and species-level random intercepts and environmental slopes partitioned among phylogenetic, trait-associated, and residual components for the final structure:

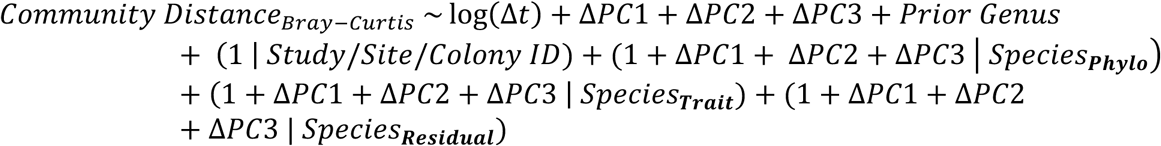

The same priors were used for shared parameters, with additional normal(0, 2) and normal(log(10), 1.5) priors placed on the beta model intercept and log precision parameter, respectively. The model was fit using the same sampling settings and convergence criteria as the ordinal model.

### Sensitivity analyses

To assess the robustness of the primary results to methodological differences and dataset scale, five additional models were fit. First, the genus-level response was reduced to a binary outcome, and observations for each colony were retained only through its first turnover event, preventing subsequent reversions or repeated events from influencing the results. Second, the genus-level ordinal model was restricted to the timeframe represented by the DIV-level model. Third, the genus-level model was repeated after excluding colonies initially observed in mixed-genus states. Fourth, prior symbiont genus was omitted as a predictor to test whether its inclusion masked environmental effects. Finally, the DIV-level beta model was repeated using weighted UniFrac rather than Bray–Curtis distances. None of these analyses materially altered the conclusions of the primary models. Full model specifications and results are provided in Supplemental File 2.

### Directional transition modeling using multi-state Markov models

The Bayesian models treat turnover severity as a directional outcome agnostic to state identity. Thus, both initial turnover events and reversions to the original symbiont state are treated equivalently, and the destination genus of a turnover event is not explicitly modeled. To address these limitations, I fit two continuous-time multi-state Markov models using the full genus-level dataset using the ‘msm’ package in R (Jackson, 2011).

The first model estimated baseline turnover and reversion rates and their dependence on environmental change (ΔPCs). Each colony was assigned to one of two states at each sampling interval: the baseline state, defined as the dominant symbiont observed at first sampling for that colony, and the novel state, defined as any departure from that baseline symbiont. This two-state formulation allowed direct estimation of transition intensities between initial turnover and reversion, and their modulation by environmental change as captured by ΔPC1, ΔPC2, and ΔPC3. Time was modeled in years elapsed since first observation for each colony. Hazard ratios for each environmental covariate were extracted from the fitted model. This model does not account for host species identity; a version with species as an additional covariate was attempted but only 12/67 species had sufficient data to fit models.

The second model estimated genus-level transition probabilities across the four major symbiont genera from the genus-level dataset (*Symbiodinium, Breviolum, Cladocopium,* and *Durusdinium*). Extending this approach to DIV-level data was impractical given the large number of potential transitions (205 observed and 5,041 theoretically possible transitions across approximately 3,000 observations). Co-dominant states were treated as censored observations, with each co-dominant genus state assigned to the set of pure states it could plausibly represent. This avoided explicitly modeling additional co-dominant states and substantially improved model convergence. Rare genera observed in <5% of colonies (e.g., *Gerakladium*) were excluded prior to fitting to reduce state space. All pairwise transitions among the four remaining states were permitted, with initial transition intensities set to 0.005. Covariates were not included because their inclusion prevented reliable convergence.

## Results

### Symbiont identity, not environmental conditions, best predicts colony-level turnover across coral species

The genus- and DIV-level Bayesian models produced consistent results. In both models, sampling interval duration and prior symbiont genus were the only fixed effects with 95% credible intervals which did not overlap zero (Tables S2-S3). Changes in environmental conditions (ΔPC1-PC3) showed little consistent effect on either genus-level turnover severity (Figure 2a; top) or DIV-level community dissimilarity (Figure 2b; top).

**Figure 2.**
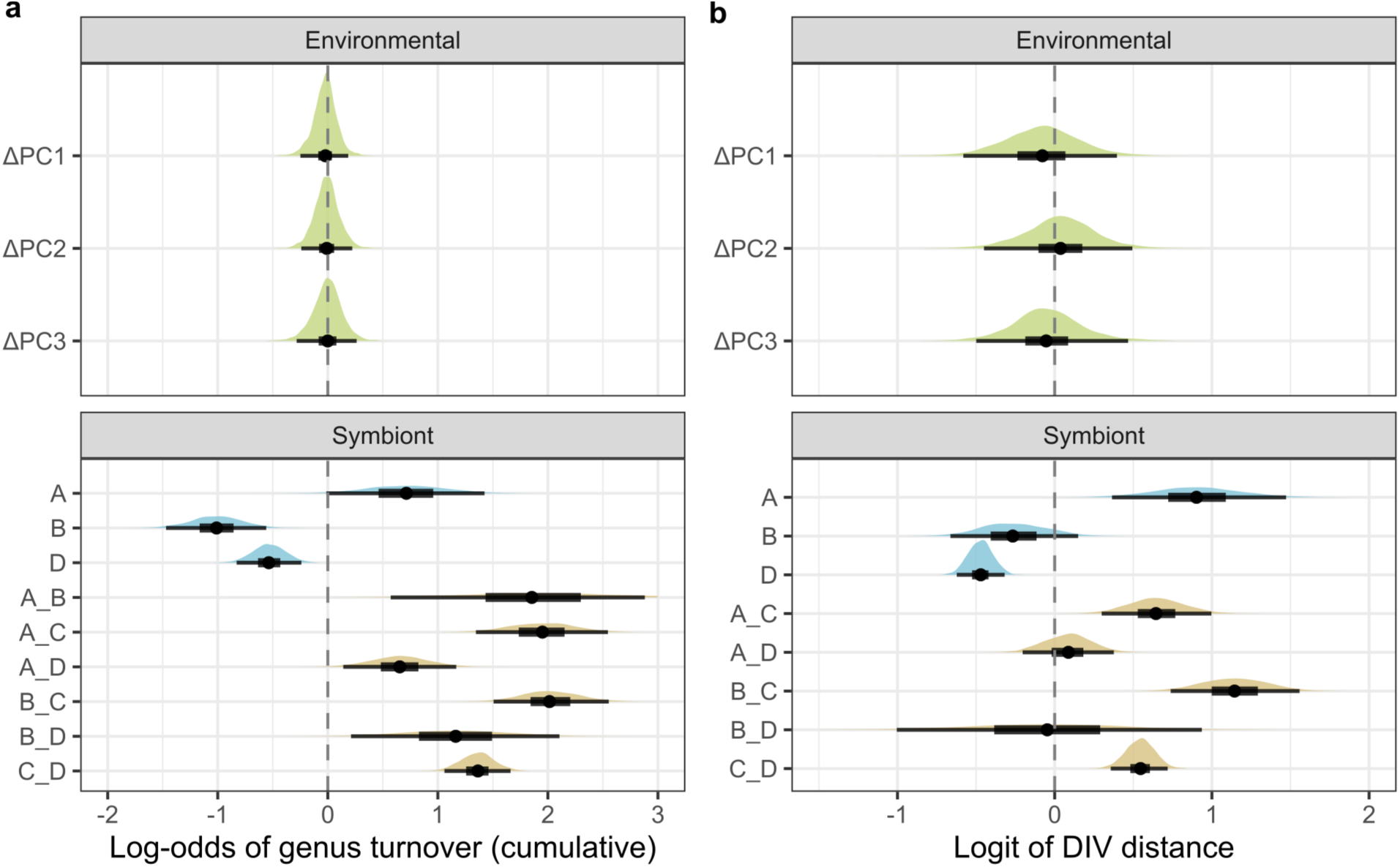
Posterior estimates of fixed effects of environmental conditions and initial symbiont identity on (a) genus-level turnover severity and (b) DIV-level community dissimilarity. Posterior distributions are shown with median estimates (points) and 50% (thick lines) and 95% (thin lines) credible intervals. Positive coefficients indicate greater turnover severity (a) or community dissimilarity (b), while negative coefficients indicate lower turnover; dashed lines indicate no effect. Environmental PCs (top) showed little consistent effect on turnover across either model (PC loadings; Figure S1). Symbiont identity (bottom) was defined by the dominant genus or combination of co-dominant genera at the beginning of each sampling interval, with *Cladocopium* (C) as the reference category. Corals initially hosting *Symbiodinium* (A), *Breviolum* (B), *Durusdinium* (D), or co-dominant genera differed in subsequent turnover. In both models, co-dominant communities (brown) generally exhibited greater turnover than communities dominated by a single genus (blue). Rare genus states were excluded from the visualization, but their posterior estimates are provided in Tables S2 and S3.

Conversely, prior symbiont genus was a strong predictor of turnover in both models. Colonies hosting co-dominant genera generally exhibited greater turnover than those dominated by a single genus, although this pattern was weaker for mixed *Symbiodinium/Breviolum* and *Durusdinium* communities. Among single-genus colonies, those hosting *Cladocopium* or *Symbiodinium* exhibited greater turnover than those hosting *Breviolum* or *Durusdinium* (Figure 2a,b; bottom).

### Host traits and phylogeny primarily determine variation in symbiont turnover across environmental gradients

Both the genus-level and DIV-level Bayesian models revealed variation in turnover among studies, reefs, and individual colonies, as well as differences among coral species in baseline turnover and sensitivity to environmental change (Figure S3; Tables S4-S5). To understand the sources of these species-level differences, I examined the estimated variance partitioning among phylogenetic similarity, trait similarity, and idiosyncratic species differences for both baseline turnover and environmental sensitivity (Figure 3).

**Figure 3.**
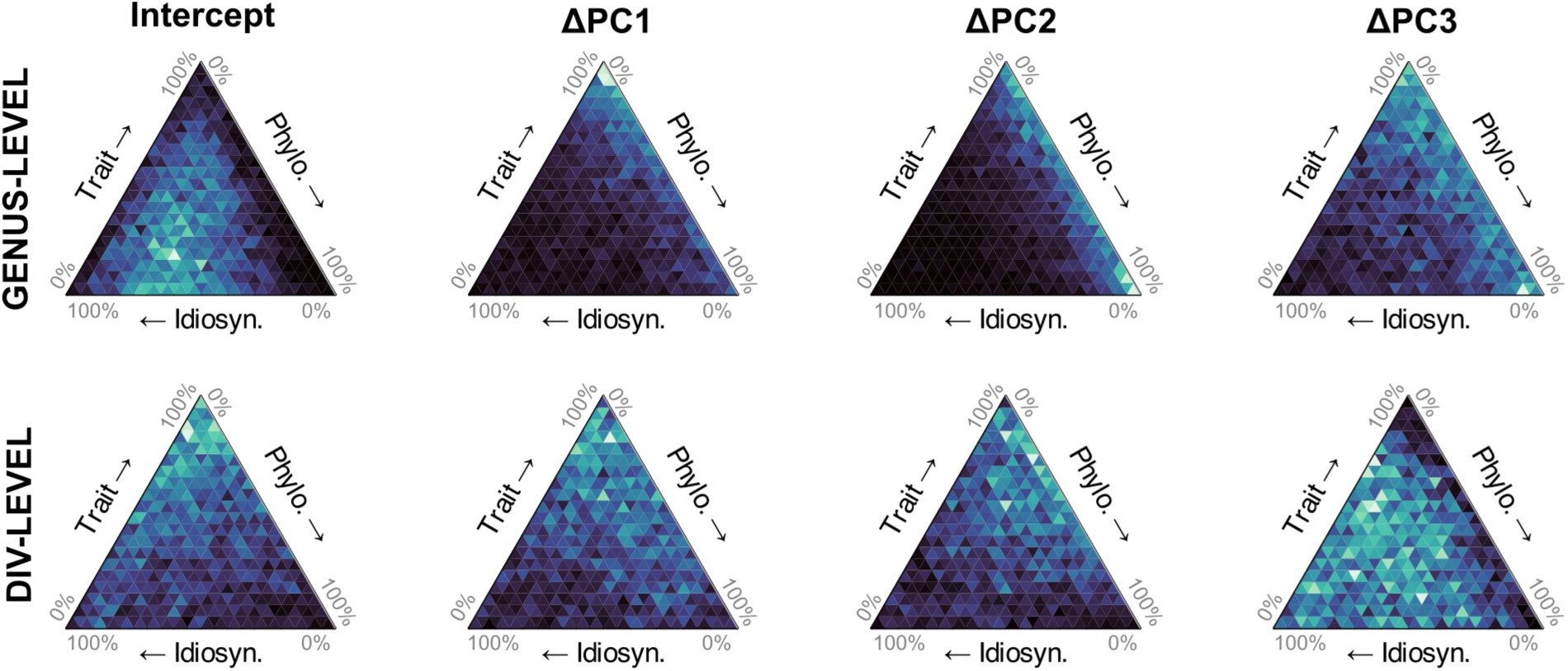
Trait and phylogenetic similarity structure species-level variation in symbiont community turnover under changing environmental conditions. Ternary density plots showing the posterior distribution of species-level variance allocated across idiosyncratic, trait-based, and phylogenetic components for baseline turnover propensity (intercept) and sensitivity to changes in each environmental principal component (ΔPC1–ΔPC3). The top and bottom rows represent genus- and DIV-level community turnover, respectively. Each axis ranges from 0% to 100% of species-level variance attributed to the component, with brighter colors indicating higher posterior density.

In both genus- and DIV-level models, sensitivity to environmental change (along ΔPC1 and ΔPC2) was better explained by phylogenetic and trait-based variation than idiosyncratic species differences. Trait similarity was the most frequently dominant component, particularly in the DIV-level model (Tables S6-S7). Baseline turnover showed a different pattern, with idiosyncratic species differences most explanatory of variation in the genus-level model and trait similarity most prominent in the DIV-level model. In the DIV-level model, no variance component consistently dominated species-level variation in sensitivity to ΔPC3. In the genus-level model, ΔPC3 showed a similar but less pronounced pattern of variance partitioning to trait and phylogenetic similarity as for ΔPC1 and ΔPC2. Species-specific turnover responses to environmental change are shown for the genus-level model in Figure 4 and the DIV-level model in Figure S4.

**Figure 4.**
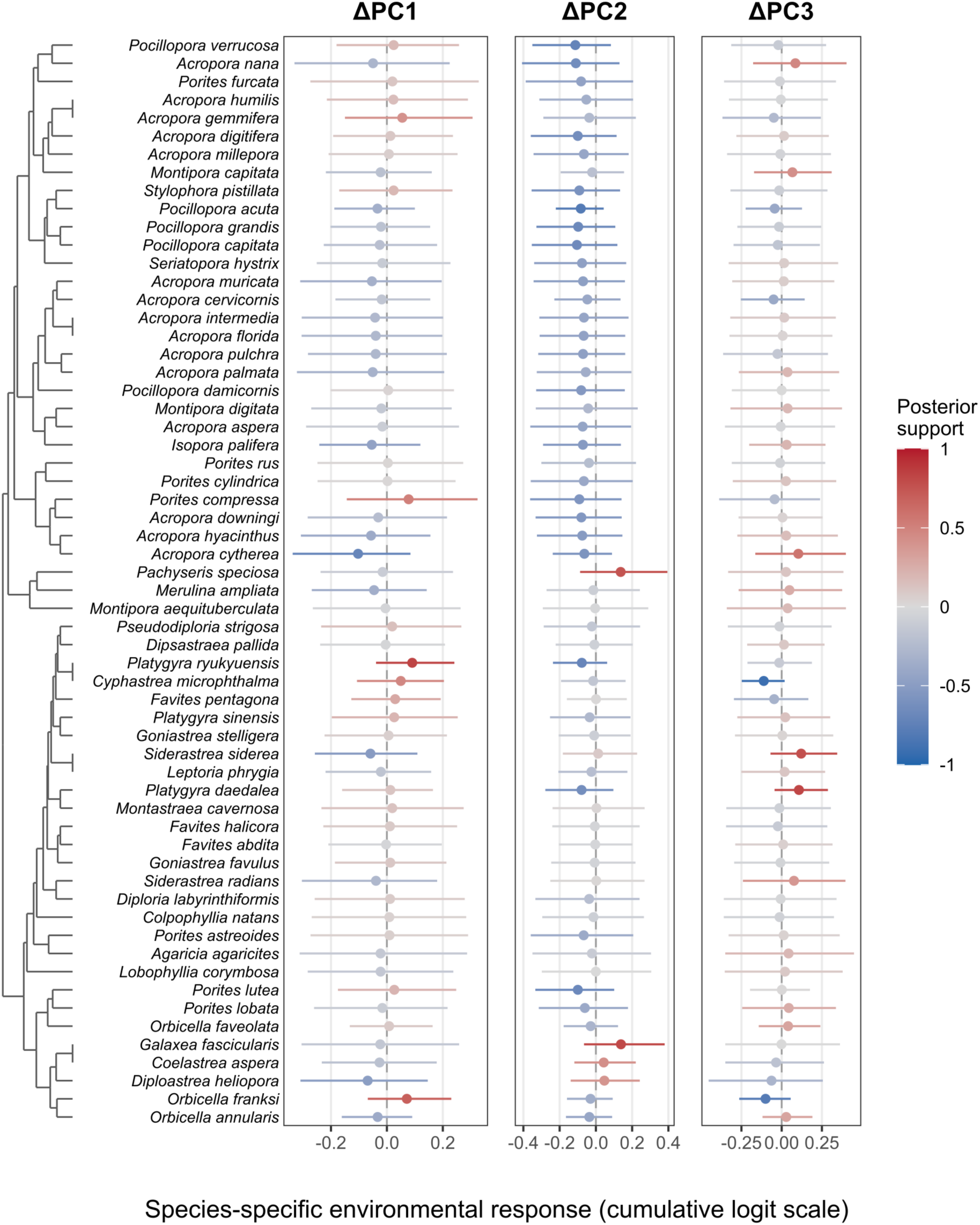
Species-specific responses in symbiont community turnover to changing environmental conditions. Posterior estimates of species-specific slopes for changes in each environmental principal component (ΔPC1–ΔPC3) from the ordinal model of genus-level symbiont turnover (DIV-level results in Figure S4). Points represent posterior median slopes, including population-level fixed effects and species-specific deviations, with horizontal bars indicating 95% credible intervals. Positive slopes indicate increased turnover with increasing environmental PC values, while negative slopes indicate decreased turnover. Colors indicate posterior support for the direction of each response, with red representing positive and blue representing negative slopes. Slopes are shown on the cumulative logit scale, with vertical reference lines indicating no effect. Species are ordered by trait similarity, as illustrated by the dendrogram on the left. A genus- and DIV-level version ordered by phylogenetic similarity can be found in Figures S5-S6.

### Annual reversion probabilities exceed those of initial genus-level turnover

The Bayesian models estimate the probability and severity of a genus-level turnover event at each sampling interval but treat initial changes and reversions equivalently and do not account for the time-dependent nature of turnover rates or the identity of the destination state. To address these limitations, I additionally fit continuous-time multi-state Markov models, which explicitly estimate transition rates between symbiont states, separate initial turnover events from reversion, and account for right censorship, where colonies with no observed turnover may simply not have been observed long enough to detect an event.

Across species, colonies initially hosting their baseline symbiont genus had a 17.26% (CI_95%_: 15.56–19.11%) probability of occupying a different genus state after one year (“Novel”). Conversely, colonies in the novel state had a 54.22% (CI_95%_: 49.57–58.91%) probability of returning to their initially observed genus after one year (“Base”; Figure S7). The hazard ratio of initial turnover decreased with ΔPC1 (CI_95%_: 0.884-0.994) and increased with ΔPC3 (CI_95%_: 1.03–1.17; Figure 5b). The same effect was not observed for ΔPC2 (CI_95%_: 0.947–1.07; Figure S7).

**Figure 5.**
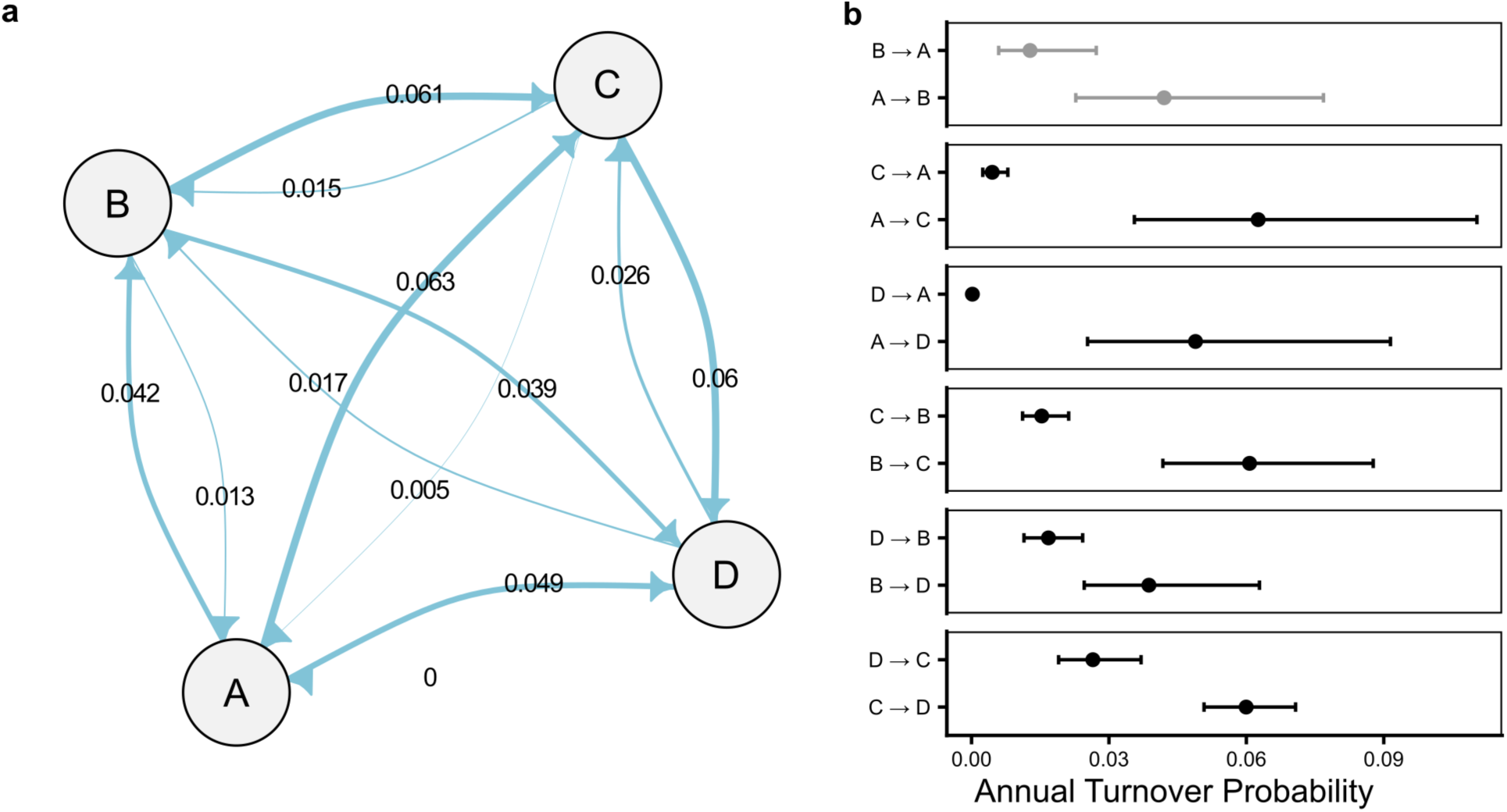
Pairwise annual transition probabilities among the four major Symbiodiniaceae genera. (a) Network diagram of mean estimated annual transition probabilities among *Symbiodinium* (A), *Breviolum* (B), *Cladocopium* (C), and *Durusdinium* (D), with arrow width proportional to transition rate. (b) Estimated annual transition rates with 95% confidence intervals for each directional pairwise transition, grouped by genus pair. Black intervals indicate pairs where confidence intervals between reciprocal transitions did not overlap.

### Symbiont turnover probabilities are not symmetrical between genera

Considering the starting and destination genus of turnover events revealed strong asymmetry in pairwise transition probabilities among Symbiodiniaceae genera (Figure 5a). Transitions from *Symbiodinium* to *Cladocopium* had the highest estimated annual probability in the dataset (0.063), though this estimate was comparable to several other transitions. Pairwise transition probabilities were frequently asymmetric, with non-overlapping 95% confidence intervals for *Symbiodinium* ↔ *Cladocopium, Symbiodinium* ↔ *Durusdinium*, *Cladocopium* ↔ *Breviolum*, *Breviolum* ↔ *Durusdinium,* and *Cladocopium* ↔ *Durusdinium* transitions. In all cases where asymmetry was detected with *Durusdinium*, transitions toward *Durusdinium* were more probable than transitions away from it (Figure 5b).

## Discussion

It is clear that environmental perturbations can drive shifts in symbiont community composition at the reef level with measurable ecological consequences (Claar et al., 2020; Glass et al., 2026). Drawing on longitudinal data spanning 67 coral species, 26 years, and 109 globally distributed reefs, I show that symbiont turnover is not a general response to environmental change. Instead, it emerges from variation in host traits and phylogeny, the resident symbiont community, and environmental conditions.

These results add nuance to expectations that warming should broadly trigger symbiont turnover which subsequently protects corals from further thermal stress (Buddemeier & Fautin, 1993; Jones et al., 2008; Silverstein et al., 2015). While the multistate model, which did not account for host species or symbiont identity, detected effects of ΔPC1 and ΔPC3 on initial turnover probability (Figure S7), parallel Bayesian analyses found no generalizable effect of either axis after accounting for species identity (Figure 2; Supplemental File 2). Together, this contrast suggests that environmental change increases turnover in only some host species, producing a detectable average signal that does not reflect a general response.

Conversely, resident symbiont genus consistently predicted turnover across coral species (Figure 2). Colonies dominated by *Breviolum* or *Durusdinium* were less likely to experience turnover than those dominated by *Cladocopium*, although multistate modeling showed that transition probability also depended on the destination genus. Higher turnover among *Cladocopium*-dominated colonies could reflect weaker competitive dominance, potentially associated with its broad host compatibility (Butler et al., 2023; though see Thornhill, LaJeunesse, et al., 2006). Alternatively, some *Cladocopium*-dominated associations may be comparatively sensitive to environmental stress (Berkelmans & van Oppen, 2006; Jones et al., 2008). Because genus states were defined using relative abundances, apparent turnover away from *Cladocopium* could reflect its loss rather than active proliferation of another genus. Nevertheless, environmental change remained unrelated to turnover when prior genus was omitted from the model (Supplemental File 2), indicating that its inclusion did not mask a general environmental effect.

Mixed-genus communities were generally transient, consistent with the predominance of a single Symbiodiniaceae lineage in adult colonies (Baker, 2003; Mieog et al., 2007). Co-dominance can arise during post-bleaching recovery (e.g., Buzzoni et al., 2026), so disturbance before monitoring began could have shaped both the communities first observed and their subsequent turnover. This possibility is particularly relevant here because many observations were collected around the 2016 mass bleaching event, and some studies may have begun while colonies were still recovering (Figure 1). At first observation, 349 of 3,027 colonies hosted co-dominant genera, with initial co-dominance appearing at highest proportion in 2011, 2017, and 2023, suggesting an increase after large-scale bleaching events (Figure S8). Because initial co-dominance may indicate that some colonies were still recovering from disturbance, I repeated the analysis after excluding these colonies. The lack of a general association between environmental change and turnover persisted (Supplemental File 2). Nonetheless, the possibility that single-genus colonies had already undergone disturbance-related turnover before monitoring began cannot be ruled out.

Transitions between dominant genera were also asymmetrical. Transitions toward *Durusdinium* were consistently more probable than transitions away from it, regardless of starting genus. Its persistence may reflect continued environmental suitability and the thermal tolerance it can confer under sustained heat (Berkelmans & van Oppen, 2006). Alternatively, *Durusdinium* may act as an opportunist that achieves dominance following community disruption and persists after conditions stabilize (Chan et al., 2026; Pettay et al., 2015). Further, this persistence does not necessarily establish a benefit to the host. For example, increased *Durusdinium* dominance has been associated with greater mortality during subsequent stress in some systems (Claar et al., 2020), but with no detectable growth consequences in others (Palacio-Castro et al., 2023). Transition asymmetry may therefore reflect environmental suitability or competitive persistence without implying a consistent host-fitness outcome.

Although environmental change had no general effect across species, species differed in their responses, and this variation was structured by trait similarity and phylogenetic relatedness (Figure 3). These axes were only moderately correlated (*Procrustes R* = 0.39, *p* = 0.002), suggesting that they captured related but partially distinct dimensions of host biology. Trait similarity most consistently structured responses to ΔPC1 and ΔPC2 at both genus and DIV resolutions. Positive ΔPC1 represented shifts toward warmer, somewhat wetter, less turbid, and less variable conditions, whereas positive ΔPC2 represented shifts toward anomalously warm, wetter, more turbid, and generally more variable conditions (Figure S1). Phylogenetic history was particularly associated with responses to ΔPC3 in the genus-level model. Positive ΔPC3 represented shifts toward higher minimum temperatures, lower thermal variability, and weaker SST anomalies—that is, consistently warm conditions rather than episodic extremes.

Coral traits have long been associated with environmental sensitivity, with slow-growing massive corals often showing greater bleaching resistance than faster-growing branching species (Darling et al., 2012; Loya et al., 2001; Marshall & Baird, 2000). Coral life-history characteristics have also been linked to algal symbiont specificity, flexibility, and thermal tolerance (Putnam et al., 2012; Swain et al., 2018). Ziegler et al. (2019) similarly proposed that broader microbial flexibility may reflect contrasting host life-history strategies. The results presented here are consistent with this hypothesis, showing that trait similarity explains a component of species-level variation in symbiont turnover across both genus- and DIV-level models. Although these patterns suggest that symbiont turnover may be associated with broader aspects of coral life history, identifying the specific traits and potential trade-offs underlying this relationship will require further investigation.

The phylogenetic signal, particularly in genus-level responses to ΔPC3, may represent a related but distinct axis of host variation. Phylogeny can capture conserved characteristics not represented in the trait matrix, including physiological tolerances, symbiont-association patterns, and other aspects of the evolutionary history of coral photosymbiosis (Gault et al., 2021; Swain et al., 2021). Because ΔPC3 contrasted persistently warm, low-variability conditions with more variable or anomalous thermal regimes, related coral lineages may respond similarly if they share unmeasured physiological characteristics or histories of exposure to comparable environments. Differences in variance partitioning between the genus- and DIV-level models raise the possibility that phylogenetically structured variation is more consequential for broad changes in dominant genus than for finer-scale community reorganization along this environmental axis.

Several limitations qualify the interpretation of these patterns. First, environmental change may influence turnover indirectly through the severity of a colony’s bleaching response rather than through environmental exposure alone. Bleaching can alter symbiont composition (e.g., Jones et al., 2008; Silverstein et al., 2015), while bleaching susceptibility is itself associated with host traits and phylogeny (Loya et al., 2001; Marshall & Baird, 2000; Swain et al., 2021). Colonies experiencing similar environmental change may therefore differ in bleaching severity and recovery, but bleaching phenotype was not consistently reported in the meta-analysis and could not be included as a predictor here. Second, reef environmental history has been linked to bleaching susceptibility (Safaie et al., 2018), Symbiodiniaceae composition and dynamics (de Souza et al., 2023; McRae et al., 2023), and the taxonomic and functional composition of coral assemblages (Darling et al., 2013; Hughes et al., 2018). Host traits, lineages, and symbiont communities may consequently covary because they have been filtered by similar environments, rather than solely because traits or shared ancestry directly regulate turnover.

Limited host and symbiont genetic resolution may further constrain inferences. Several well-represented taxa belong to cryptic species complexes, including *Porites* and *Pocillopora*, within which distinct host lineages associate with different Symbiodiniaceae communities (Grupstra et al., 2024; Johnston et al., 2022; Scott et al., 2025). Pooling these lineages may obscure lineage-specific dynamics and overestimate flexibility at the nominal species level. Initial symbiont composition can vary with host genotype within a species and may influence which background symbionts proliferate during stress and recovery by shaping their starting abundances (Baker, 2003; Bay et al., 2016; Quigley et al., 2019). Finally, although DIV profiles resolve finer community differences than genus-level classifications, they cannot resolve whether the same clonal symbiont strains persisted through time. Given the prevalence of fine-scale host-symbiont specificity (e.g., Hume et al., 2020), future studies pairing genome-scale host and symbiont characterization with repeated community sampling will help determine how these differences contribute to turnover.

The present synthesis documents patterns of turnover across species and environments, but their fitness consequences remain largely unresolved. Single-species experiments indicate that symbiont community shifts can increase thermal tolerance or other components of fitness while imposing physiological costs, with outcomes that depend on the host–symbiont combination and environmental context (Cunning et al., 2015; Olivares-Cordero et al., 2025). Determining whether turnover represents acclimatization, dysbiosis, or a largely neutral change will require longitudinal measurements of host growth, survival, reproduction, and subsequent stress tolerance. The phylogenetic and trait axes characterized here provide a scaffold for testing whether these consequences vary predictably across coral life histories, complementing evidence that both axes structure symbiont associations and bleaching susceptibility more broadly (Swain et al., 2021; Williams & Patterson, 2020; Zarate et al., 2024).

## Conclusion

Symbiont turnover across scleractinian corals is not a universal response to changing conditions, but an emergent outcome shaped by host traits, phylogenetic history, and symbiont ecology. These results challenge the assumption that symbiotic flexibility should generally be expected under changing environmental conditions. Decades of observational data have documented variation in turnover across reef systems worldwide, yet the drivers of that variation have remained unclear; the results presented here offer a framework for understanding when, and in which species, turnover is most likely to occur.

## Acknowledgements

I thank Christopher Peterson for opening my eyes to the breadth of possibilities in Bayesian modeling and, in particular, for his guidance on prior determination for this analysis. I also thank Charles Mitchell, Schyler Ellsworth, Sherlynette Castro, Kristina Black, and Dom Gallery for their valuable feedback at various stages of analysis and writing.

## Data Availability

All raw data, analysis and processing scripts, and intermediate/final data products can be found at https://github.com/cb-scott/longitudinal_symbiodiniaceae. Final versions will be archived on Zenodo upon paper acceptance.

