## Supplemental File 1 for "Within-host algal symbiont turnover is not a general response to environmental change in reef corals"

### Appendix S1: Supplemental Figures and Tables Supporting the Core Models

Within-host algal symbiont turnover is not a general response to environmental change in reef corals

Carly B. Scott

#### Supplemental Figures

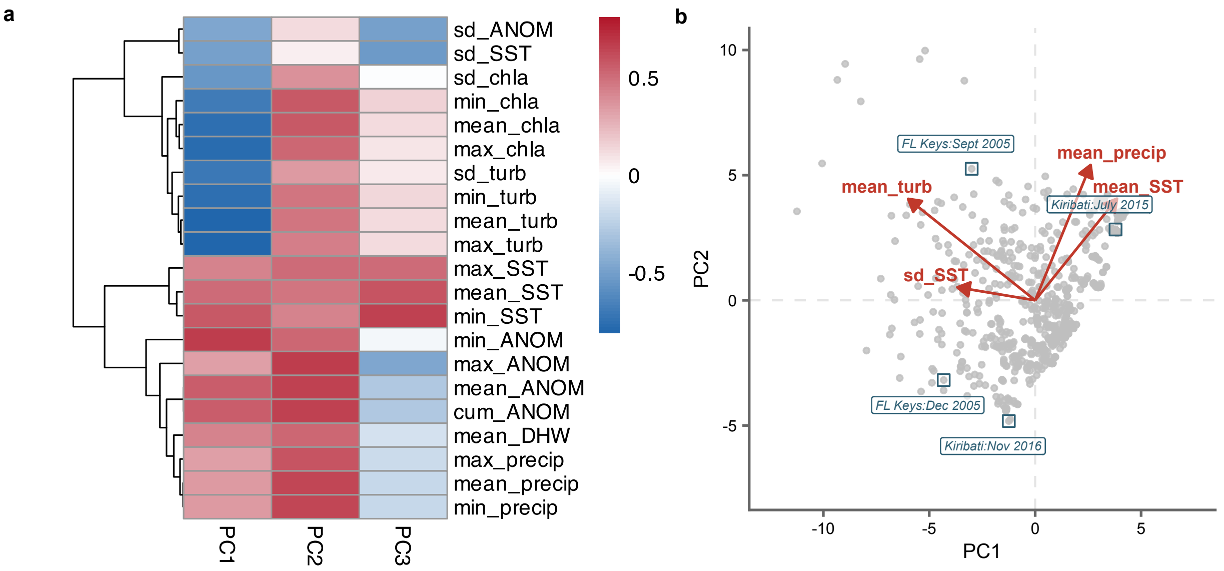


**Supplemental Figure S1.** (A) Loadings on environmental variables with environmental PC1, PC2, and PC3. Prefixes summarize each variable over the month preceding observation: min, minimum; max, maximum; sd, standard deviation; cum, cumulative sum. Environmental variables are SST, sea surface temperature; ANOM, SST anomaly; chla, chlorophyll a; turb, turbidity measured as kd490; precip, estimated precipitation; and DHW, degree heating weeks. (B) Environmental PCA ordination of observations included in the study. Red arrows show representative variable loadings on PC1 and PC2, and annotated points identify examples of major bleaching and recovery observations represented in the dataset.

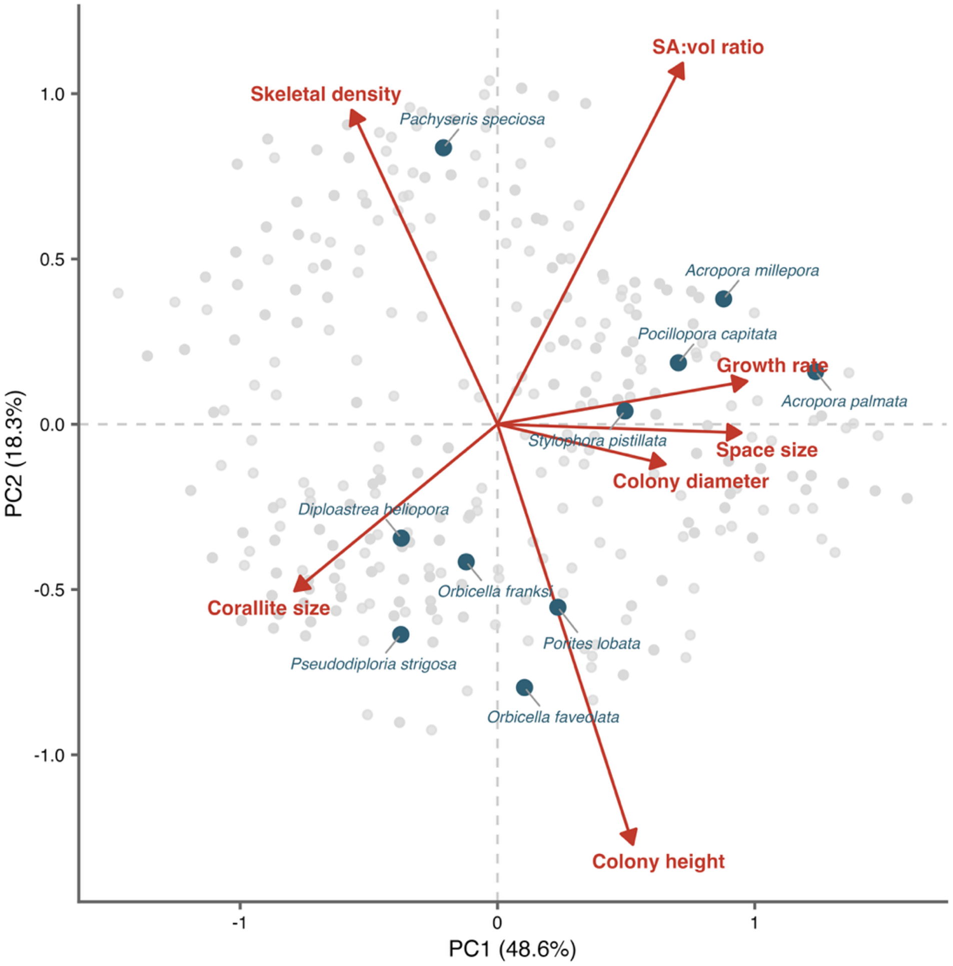


**Supplemental Figure S2.** Principal-components ordination of coral functional traits compiled from McWilliam et al. (2018). Each point represents a coral species used to construct the trait covariance matrix. Annotated species illustrate contrasting growth strategies represented in the analysis, and arrows show the loadings of the included traits on the first two principal components.

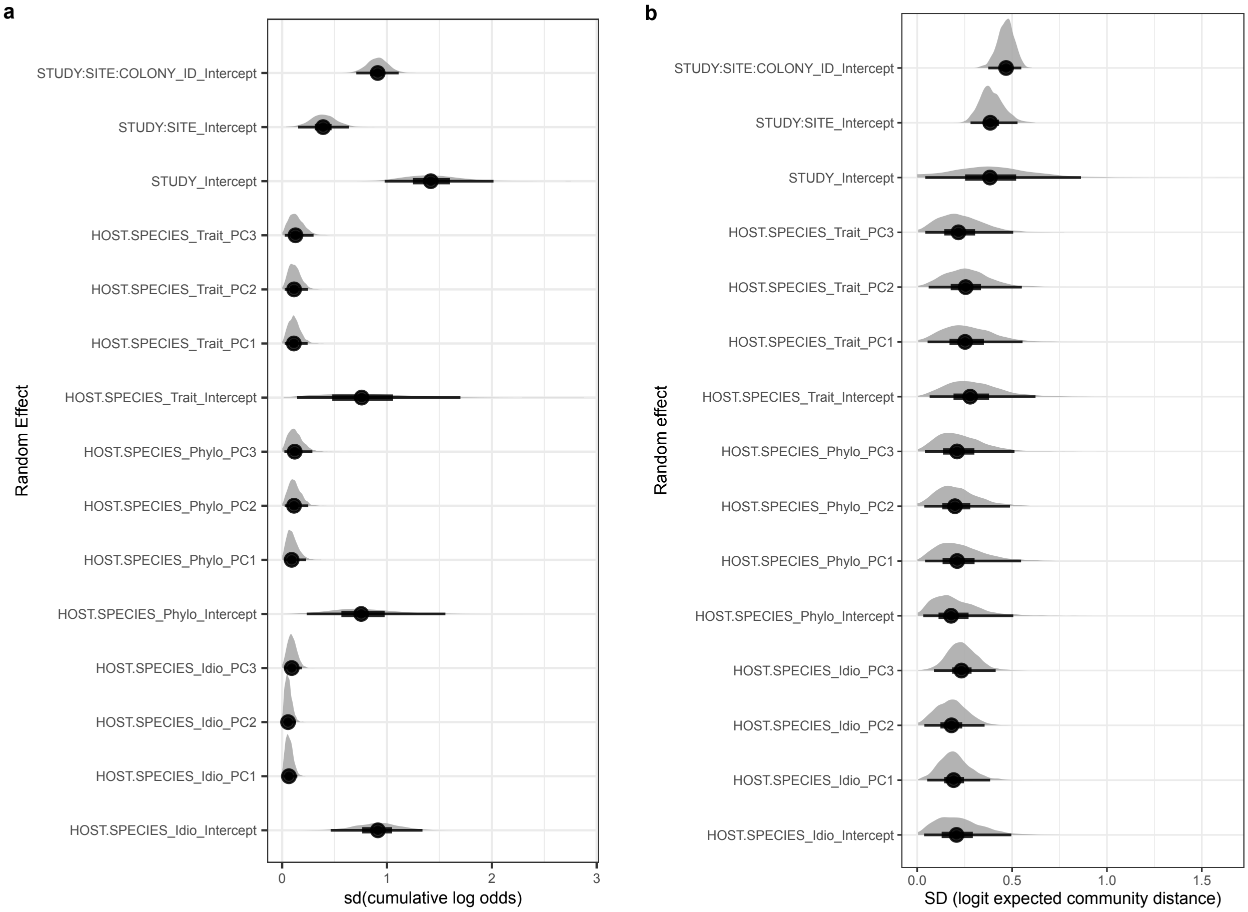


**Supplemental Figure S3.** Posterior distributions of random-effect standard deviations from the two primary Bayesian hierarchical models. (a) Genus-level ordinal model. (b) DIV-level Bray-Curtis beta model. Distributions summarize uncertainty in among-group variation for study, site, and colony intercepts and for species-level intercepts and environmental slopes partitioned among phylogenetic, trait-associated, and idiosyncratic components. Points show posterior medians and shaded densities show the posterior distributions.

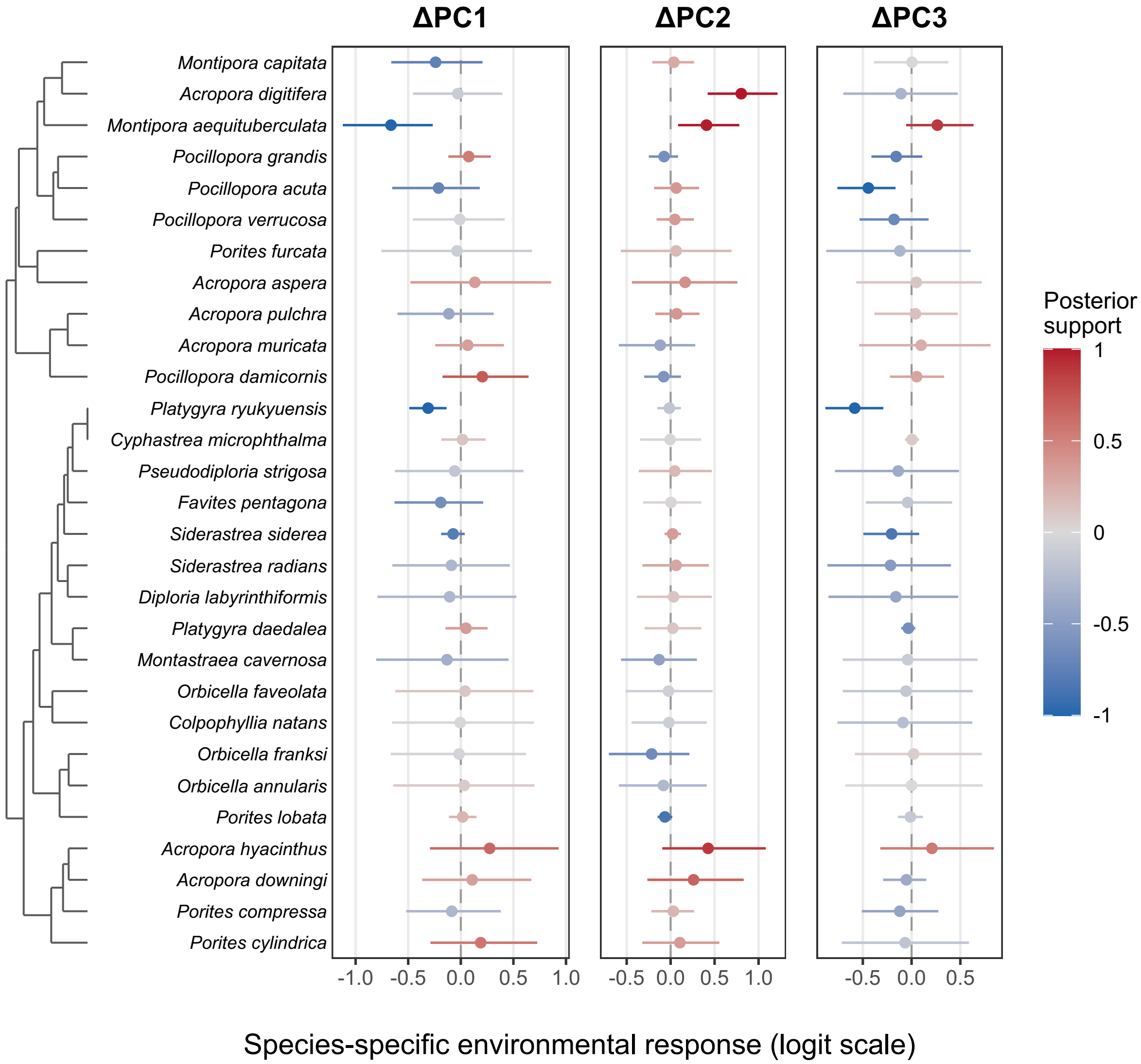


**Supplemental Figure S4.** Species-specific environmental responses from the primary DIV-level Bray-Curtis model. Points are posterior median total slopes, combining the population-level fixed effect with each species-specific deviation, and horizontal intervals are 95% credible intervals. Positive values indicate increasing expected DIV-level community dissimilarity as the corresponding environmental principal component increases; negative values indicate decreasing dissimilarity. Species are ordered by trait similarity, shown by the dendrogram at left. See Supplemental Figure S6 for ordering by phylogenetic similarity.

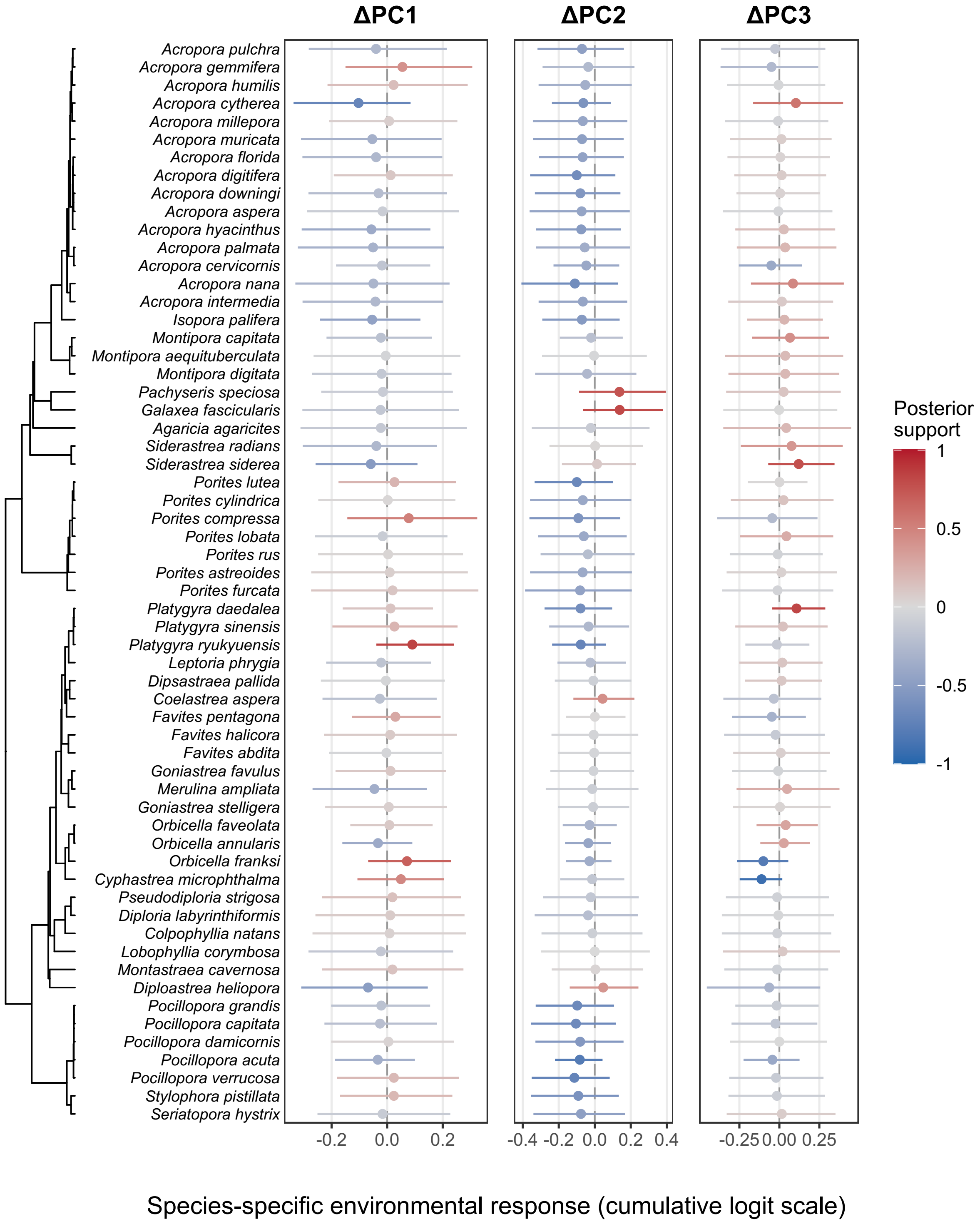


**Supplemental Figure S5.** Species-specific environmental responses from the genus-level ordinal model, ordered by phylogenetic similarity. Points are posterior median total slopes, combining the population-level fixed effect with each species-specific deviation, and horizontal intervals are 95% credible intervals.

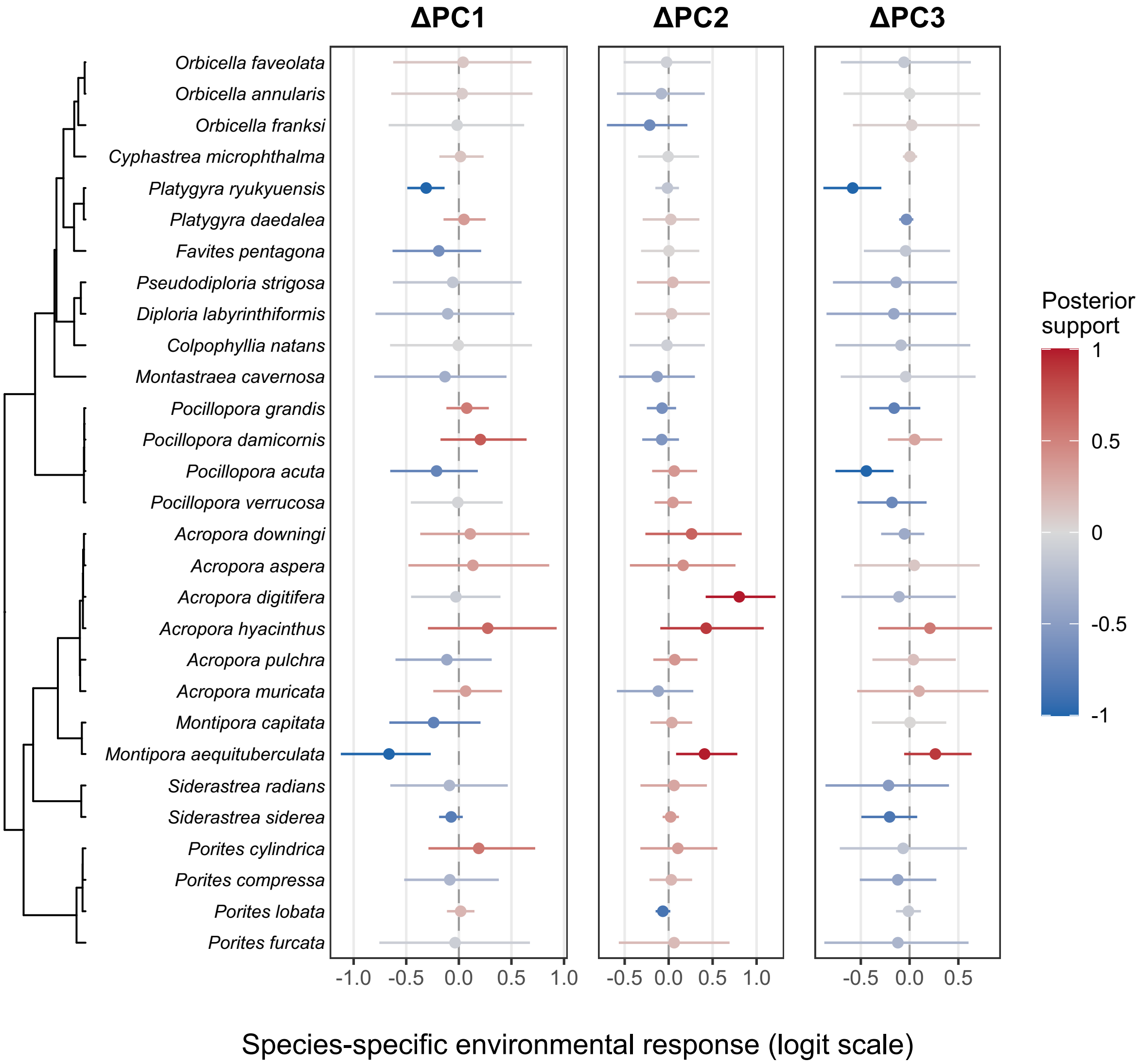


**Supplemental Figure S6.** Species-specific environmental responses from the primary DIV-level Bray-Curtis model. Points are posterior median total slopes, combining the population-level fixed effect with each species-specific deviation, and horizontal intervals are 95% credible intervals. Positive values indicate increasing expected DIV-level community dissimilarity as the corresponding environmental principal component increases; negative values indicate decreasing dissimilarity. Species are ordered by phylogenetic similarity, shown by the dendrogram at left.

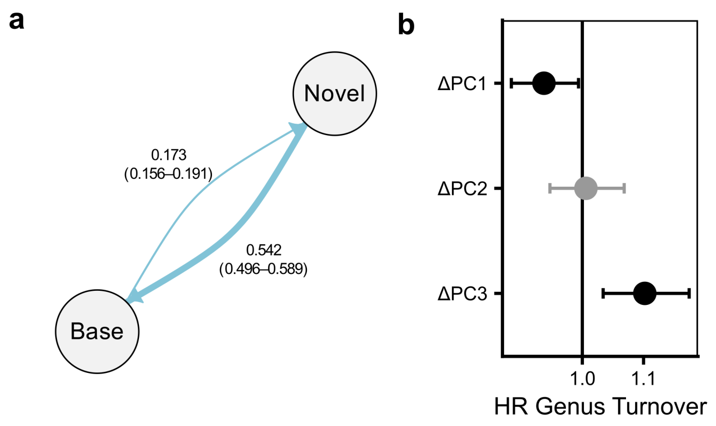


**Supplemental Figure S7. Multistate modeling of annual symbiont initial turnover and reversion rates at the genus level.** (a) Estimated annual transition probability across all species, showing the probability of an initial turnover event ("Novel", 0.173 per year) and reversion to the baseline symbiont state ("Base", 0.542 per year) for colonies that had switched. Baseline symbiont state was defined as the dominant symbiont genus observed at first sampling for each colony. (b) Hazard ratios for the effect of environmental principal components on initial turnover probability, with 95% confidence intervals. A hazard ratio greater than one indicates increased turnover probability and a hazard ratio of one (vertical line) indicates no effect.

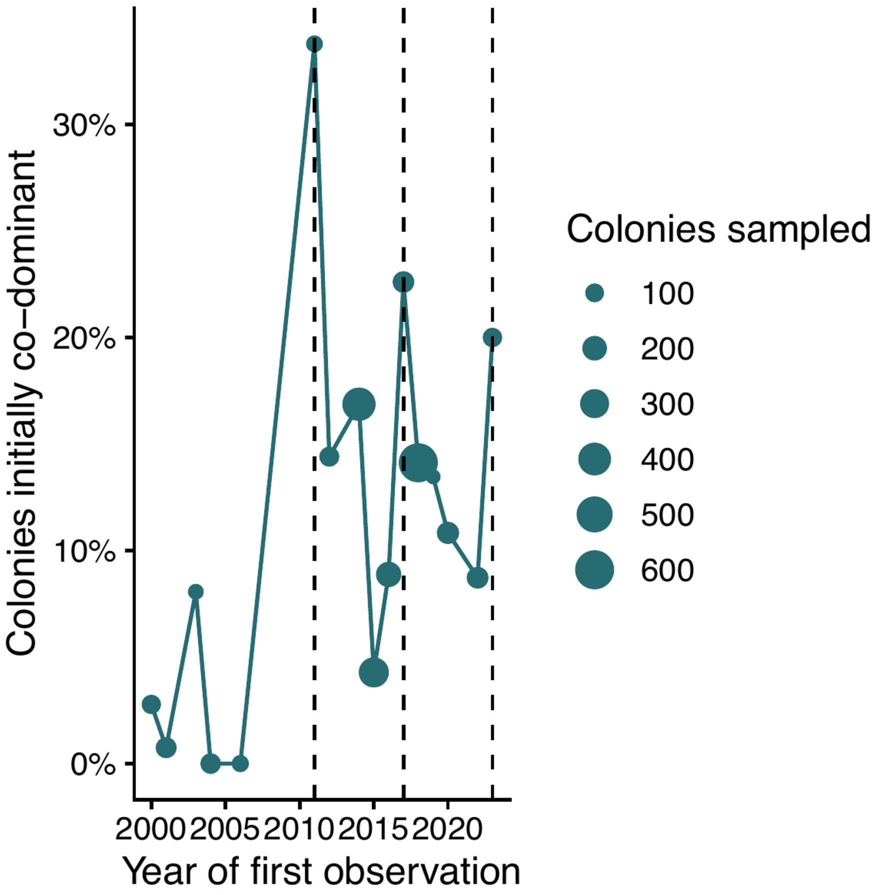


**Supplemental Figure S8.** Proportion of colonies first observed in a mixed-genus state by year of first observation. Points are scaled by the number of colonies first observed in each year; only years with more than 50 colonies are shown. Dashed vertical lines mark 2011, 2017, and 2023, which were preceded by large scale bleaching events. Across all years, 349 of 3,027 colonies (11.5%) were initially observed in mixed-genus states.

#### Supplemental Tables

**Table S1.** Boolean search strings used to identify studies for inclusion in the longitudinal synthesis.

| ***Database*** | **Search string** |
|---|---|
| *PubMed* | ("Scleractinia" OR "scleractinian" OR "hard coral" OR "reef-building coral" OR "reef coral" OR "coral coloni*") AND ("Symbiodiniaceae" OR "Symbiodinium" OR "zooxanthell*" OR "endosymbiont*") AND ("symbiont shuffling" OR "symbiont switching" OR "shuffling" OR "switching" OR "tagged colon*" OR "marked colon*" OR "longitudinal" OR "time series" OR "repeated sampling" OR "repeated measures" OR "temporal dynamics" OR "temporal variation" OR "temporal change" OR "over time" OR "same coloni*" OR "individual coloni*" OR "time point*" OR "clade" OR "ITS2" OR "symbiont type" OR "symbiont community" OR "algal symbiont*" OR "Cladocopium" OR "Durusdinium" OR "Breviolum" OR "community composition" OR "community change") |
| *Google Scholar* | coral "tagged colonies" Symbiodiniaceae clade "repeated sampling" OR "time series" OR "seasonal" |
| *Google Scholar* | "symbiont switching" OR “symbiont dynamics” AND "coral" |

**Supplemental Table S2.** Posterior medians and 95% credible intervals for fixed effects from the primary genus-level ordinal model. Fixed effects are on the cumulative-logit scale; positive coefficients indicate greater turnover severity. Cladocopium is the reference prior-symbiont state.

| ***Term*** | ***Median*** | ***2.5%*** | ***97.5%*** |
|---|---|---|---|
| *Log elapsed time* | 0.334 | 0.169 | 0.508 |
| *ΔPC1* | -0.021 | -0.249 | 0.186 |
| *ΔPC2* | -0.009 | -0.241 | 0.222 |
| *ΔPC3* | -0.001 | -0.283 | 0.262 |
| *Prior symbiont state: Symbiodinium* | 0.714 | -0.014 | 1.424 |
| *Prior symbiont state: Symbiodinium + Breviolum* | 1.903 | 0.587 | 3.269 |
| *Prior symbiont state: Symbiodinium + Cladocopium* | 1.950 | 1.346 | 2.546 |
| *Prior symbiont state: Symbiodinium + Cladocopium + Durusdinium* | 1.173 | 0.180 | 2.163 |
| *Prior symbiont state: Symbiodinium + Cladocopium + Gerakladium* | 1.506 | -0.029 | 2.992 |
| *Prior symbiont state: Symbiodinium + Durusdinium* | 0.654 | 0.141 | 1.168 |
| *Prior symbiont state: Breviolum* | -1.011 | -1.471 | -0.560 |
| *Prior symbiont state: Breviolum + Cladocopium* | 2.015 | 1.506 | 2.553 |
| *Prior symbiont state: Breviolum + Cladocopium + Durusdinium* | 0.671 | -0.949 | 2.226 |
| *Prior symbiont state: Breviolum + Durusdinium* | 1.162 | 0.211 | 2.105 |
| *Prior symbiont state: Cladocopium + Durusdinium* | 1.364 | 1.060 | 1.660 |
| *Prior symbiont state: Cladocopium + Gerakladium* | 0.531 | -1.314 | 2.291 |
| *Prior symbiont state: Durusdinium* | -0.536 | -0.828 | -0.240 |
| *Prior symbiont state: Durusdinium + Gerakladium* | 0.229 | -1.507 | 2.000 |
| *Intercept [1]: no genus turnover vs. codominance or major turnover* | 2.369 | 1.312 | 3.351 |
| *Intercept [2]: no turnover or codominance vs. major turnover* | 4.050 | 2.989 | 5.042 |

**Supplemental Table S3.** Posterior medians and 95% credible intervals for fixed effects from the DIV-level Bray-Curtis beta model. Fixed effects are on the logit scale of expected Bray-Curtis dissimilarity; positive coefficients indicate greater community dissimilarity. *Cladocopium* is the reference prior-symbiont state.

| *Term* | *Median* | *2.5%* | *97.5%* |
|---|---|---|---|
| *Log elapsed time* | 0.096 | 0.015 | 0.178 |
| *ΔPC1* | -0.078 | -0.581 | 0.396 |
| *ΔPC2* | 0.037 | -0.450 | 0.494 |
| *ΔPC3* | -0.054 | -0.499 | 0.467 |
| *Prior symbiont state: Symbiodinium* | 0.902 | 0.365 | 1.472 |
| *Prior symbiont state: Symbiodinium + Cladocopium* | 0.644 | 0.299 | 0.997 |
| *Prior symbiont state: Symbiodinium + Cladocopium + Durusdinium* | -0.131 | -1.658 | 1.354 |
| *Prior symbiont state: Symbiodinium + Durusdinium* | 0.087 | -0.204 | 0.376 |
| *Prior symbiont state: Breviolum* | -0.267 | -0.663 | 0.149 |
| *Prior symbiont state: Breviolum + Cladocopium* | 1.144 | 0.738 | 1.558 |
| *Prior symbiont state: Breviolum + Durusdinium* | -0.049 | -1.030 | 0.936 |
| *Prior symbiont state: Cladocopium + Durusdinium* | 0.546 | 0.357 | 0.719 |
| *Prior symbiont state: Durusdinium* | -0.470 | -0.623 | -0.318 |
| *Beta mean intercept* | -0.991 | -1.580 | -0.429 |
| *Beta precision (φ)* | 2.997 | 2.734 | 3.278 |

**Table S4.** Posterior medians and 95% credible intervals for random-effect standard deviations from the primary genus-level ordinal model. Values quantify among-group variation in random intercepts and species-level slopes on the cumulative-logit scale; larger values indicate greater variation.

| ***Random effect*** | ***Median*** | ***2.5%*** | ***97.5%*** |
|---|---|---|---|
| *Species phylogenetic component: intercept* | 0.754 | 0.235 | 1.557 |
| *Species phylogenetic component: ΔPC1 slope* | 0.091 | 0.017 | 0.228 |
| *Species phylogenetic component: ΔPC2 slope* | 0.113 | 0.024 | 0.248 |
| *Species phylogenetic component: ΔPC3 slope* | 0.119 | 0.021 | 0.288 |
| *Species idiosyncratic component: intercept* | 0.913 | 0.463 | 1.338 |
| *Species idiosyncratic component: ΔPC1 slope* | 0.064 | 0.012 | 0.145 |
| *Species idiosyncratic component: ΔPC2 slope* | 0.056 | 0.011 | 0.133 |
| *Species idiosyncratic component: ΔPC3 slope* | 0.091 | 0.019 | 0.189 |
| *Species trait component: intercept* | 0.758 | 0.143 | 1.700 |
| *Species trait component: ΔPC1 slope* | 0.112 | 0.025 | 0.243 |
| *Species trait component: ΔPC2 slope* | 0.114 | 0.025 | 0.247 |
| *Species trait component: ΔPC3 slope* | 0.127 | 0.025 | 0.299 |
| *Study intercept* | 1.417 | 0.978 | 2.016 |
| *Site-within-study intercept* | 0.391 | 0.153 | 0.638 |
| *Colony-within-site intercept* | 0.911 | 0.707 | 1.111 |

**Table S5.** Posterior medians and 95% credible intervals for random-effect standard deviations from the primary genus-level ordinal model. Values quantify among-group variation in random intercepts and species-level slopes on the cumulative-logit scale; larger values indicate greater variation.

| ***Random effect*** | ***Median*** | ***2.5%*** | ***97.5%*** |
|---|---|---|---|
| *Species phylogenetic component: intercept* | 0.166 | 0.033 | 0.470 |
| *Species phylogenetic component: ΔPC1 slope* | 0.218 | 0.039 | 0.552 |
| *Species phylogenetic component: ΔPC2 slope* | 0.200 | 0.041 | 0.474 |
| *Species phylogenetic component: ΔPC3 slope* | 0.202 | 0.039 | 0.529 |
| *Species idiosyncratic component: intercept* | 0.188 | 0.040 | 0.446 |
| *Species idiosyncratic component: ΔPC1 slope* | 0.196 | 0.053 | 0.391 |
| *Species idiosyncratic component: ΔPC2 slope* | 0.176 | 0.038 | 0.353 |
| *Species idiosyncratic component: ΔPC3 slope* | 0.234 | 0.096 | 0.427 |
| *Species trait component: intercept* | 0.233 | 0.053 | 0.566 |
| *Species trait component: ΔPC1 slope* | 0.265 | 0.063 | 0.586 |
| *Species trait component: ΔPC2 slope* | 0.242 | 0.054 | 0.513 |
| *Species trait component: ΔPC3 slope* | 0.213 | 0.041 | 0.516 |
| *Study intercept* | 0.367 | 0.038 | 0.802 |
| *Site-within-study intercept* | 0.397 | 0.286 | 0.541 |
| *Colony-within-site intercept* | 0.462 | 0.363 | 0.544 |

**Table S6.** Posterior allocation of species-level variance among trait-based, phylogenetic, and idiosyncratic components in the genus-level models. For each model coefficient, "Leading" is the percentage of posterior draws in which a component accounted for more variance than either other component. Median share when leading is the median percentage of species-level variance assigned to that component among those draws. The intercept represents baseline turnover propensity in the genus-level model.

| | Traits | | Phylogeny | | Idiosyncratic | |
|---|---|---|---|---|---|---|
| | **Leading (%)** | **Median share when leading (%)** | **Leading (%)** | **Median share when leading (%)** | **Leading (%)** | **Median share when leading (%)** |
| **Intercept** | 30.3 | 54.3 | 23.1 | 51.8 | 46.6 | 55.9 |
| **ΔPC1** | 52.7 | 66.5 | 34.0 | 63.0 | 13.3 | 56.0 |
| **ΔPC2** | 47.9 | 66.4 | 46.9 | 67.4 | 5.2 | 49.6 |
| **ΔPC3** | 44.0 | 63.5 | 37.5 | 63.0 | 18.5 | 55.0 |

**Table S7.** Posterior allocation of species-level variance among trait-based, phylogenetic, and idiosyncratic components in the Bray-Curtis DIV-level model. For each model coefficient, "Leading" is the percentage of posterior draws in which a component accounted for more variance than either other component. Median share when leading is the median percentage of species-level variance assigned to that component among those draws. The intercept represents baseline Bray-Curtis dissimilarity in the DIV-level model.

| | Traits | | Phylogeny | | Idiosyncratic | |
|---|---|---|---|---|---|---|
| | **Leading (%)** | **Median share when leading (%)** | **Leading (%)** | **Median share when leading (%)** | **Leading (%)** | **Median share when leading (%)** |
| **Intercept** | 48.9 | 63.4 | 20.9 | 55.2 | 30.2 | 58.4 |
| **ΔPC1** | 48.1 | 61.9 | 31.1 | 58.4 | 20.8 | 56.3 |
| **ΔPC2** | 48.4 | 60.7 | 28.3 | 56.4 | 23.4 | 55.5 |
| **ΔPC3** | 32.2 | 55.3 | 28.9 | 54.8 | 38.9 | 58.0 |
