## Supplemental File 2 for "Within-host algal symbiont turnover is not a general response to environmental change in reef corals"

### Appendix S2: Supplemental Methods and Results for Sensitivity Analyses

Within-host algal symbiont turnover is not a general response to environmental change in reef corals

Carly B. Scott

#### Supplemental Methods

Five analyses evaluated whether the conclusions of the primary genus-level ordinal and DIV-level Bray-Curtis models depended on response definition, temporal coverage, initial community composition, inclusion of prior genus, or the distance metric used for DIV-level turnover. Unless noted otherwise, the analyses used the same environmental predictors and the same partition of species-level intercepts and slopes among phylogenetic, trait-associated, and idiosyncratic components as the corresponding primary model.

*Reduced-timeframe genus-level ordinal model*

The genus-level ordinal analysis was repeated after restricting observations to 30 August 2012 through 28 April 2024, the period represented in the DIV-level dataset. The response categories, cumulative-logit likelihood, fixed effects, nested study-site-colony intercepts, species-level variance partition, priors, and posterior sampling settings were otherwise unchanged from the primary ordinal analysis. This comparison tested whether the broader temporal coverage of the main genus-level dataset explained differences between the genus- and DIV-level results.

*First-turnover Bernoulli model*

The genus-level ordinal turnover response was collapsed to a binary outcome indicating no detected genus-level turnover (0) or any change in dominant or co-dominant genus state (1). For each colony, observations were retained only through the first modeled turnover event; later observations were excluded so that reversions or repeated turnover could not influence the result. A Bernoulli likelihood with a logit link replaced the cumulative ordinal likelihood. Fixed effects for elapsed time, changes in environmental PC1-PC3, and prior symbiont state were retained, as were the nested study, site, and colony intercepts and the phylogenetic, trait-associated, and idiosyncratic species-level intercepts and slopes.

*Weighted-UniFrac distance assignment and model*

At the DIV level, normalized weighted-UniFrac distances were calculated between sequential samples to test whether turnover among more evolutionarily divergent symbiont lineages differed from total DIV-level turnover. A phylogenetic tree was constructed from sequence similarities among DIVs represented in the SymPortal reference database. Direct weighted-UniFrac distance was not assigned when any DIV absent from the reference tree represented at least 10% of a community or when unmatched DIVs collectively represented more than 50% of a community; these criteria affected 375 of 2,753 comparisons.

For comparisons without sufficient DIV-level phylogenetic coverage, a calibrated hybrid estimate combined a genus-level phylogenetic proxy with within-genus Bray-Curtis turnover. Genus relative abundance was distributed equally across the sequences from that genus represented in the reference tree, and weighted UniFrac was calculated from the reconstructed communities. Within-genus turnover was then incorporated using a scaling parameter calibrated against directly calculated weighted-UniFrac distances across studies. If neither direct nor hybrid estimation was possible, comparisons with Bray-Curtis distance below 0.10 were assigned a weighted-UniFrac distance of zero and the remaining comparisons were excluded.

The weighted-UniFrac distances were analyzed with the same hierarchical beta-regression structure used for the primary Bray-Curtis model, including fixed effects for elapsed time and environmental change, nested study-site-colony intercepts, and species-level intercepts and environmental slopes partitioned among phylogenetic, trait-associated, and idiosyncratic components.

*Initially single-genus colonies*

The primary genus-level ordinal model was repeated after excluding colonies whose first observation contained a mixed-genus state. Initial state was assigned from the full observation-level dataset before sequential intervals were constructed. The model formula, likelihood, covariance structures, priors, and sampling settings were otherwise unchanged. This analysis tested whether transient mixed communities present at the first observation drove the primary results.

*Model without prior genus*

The primary genus-level ordinal model was repeated after removing prior symbiont genus from the fixed-effect design matrix while retaining all otherwise eligible observations. Fixed effects for elapsed time and changes in environmental PC1-PC3 remained in the model, and the likelihood, nested sampling structure, covariance matrices, priors, sampling settings, and species-level variance partition were unchanged. This analysis tested whether adjustment for prior genus masked a general association between environmental change and turnover.

#### Supplemental Figures

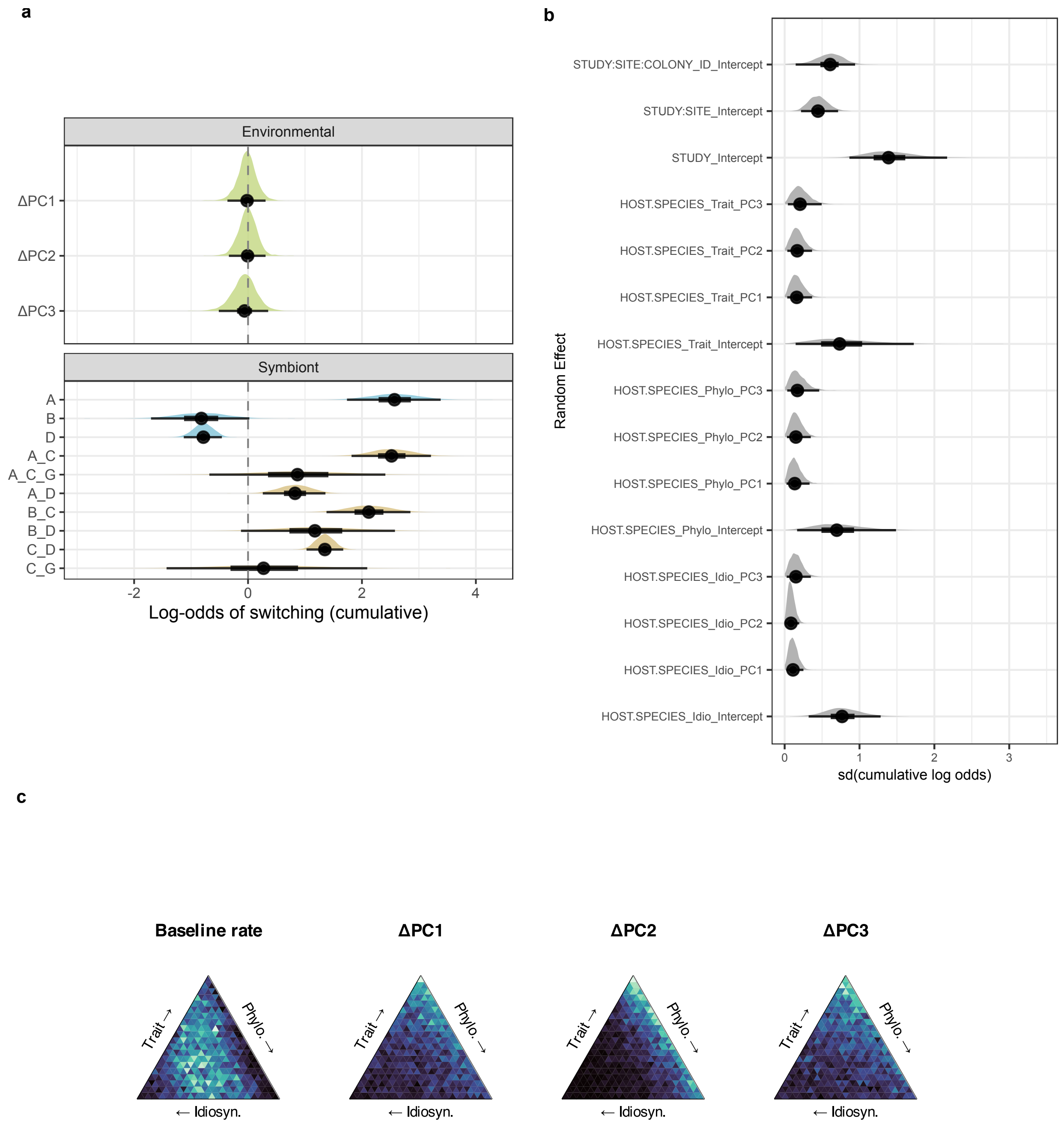


**Figure S2.1** Reduced-timeframe genus-level ordinal sensitivity analysis. The ordinal model was restricted to observations from 30 August 2012 through 28 April 2024, matching the temporal extent of the DIV dataset. (a) Posterior distributions of fixed effects. (b) Posterior distributions of random-effect standard deviations. (c) Posterior allocation of species-level variance among phylogenetic, trait-associated, and idiosyncratic components for baseline turnover and sensitivity to ΔPC1-ΔPC3.

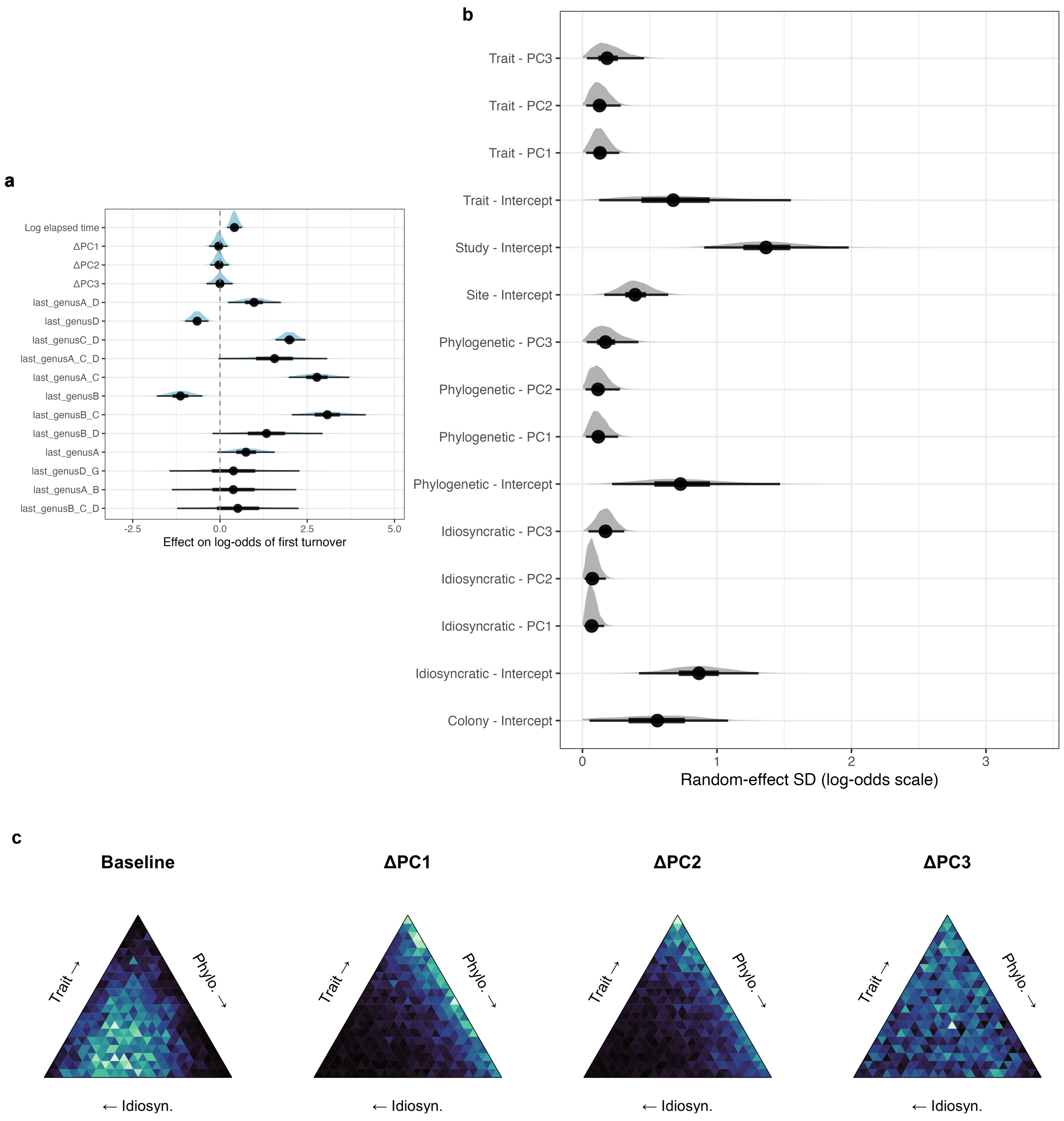


**Figure S2.2** First-turnover Bernoulli sensitivity analysis. Each colony contributed observations only through its first modeled turnover event. (a) Posterior distributions of fixed effects. (b) Posterior distributions of random-effect standard deviations. (c) Posterior allocation of species-level variance among phylogenetic, trait-associated, and idiosyncratic components for baseline turnover and sensitivity to ΔPC1-ΔPC3.

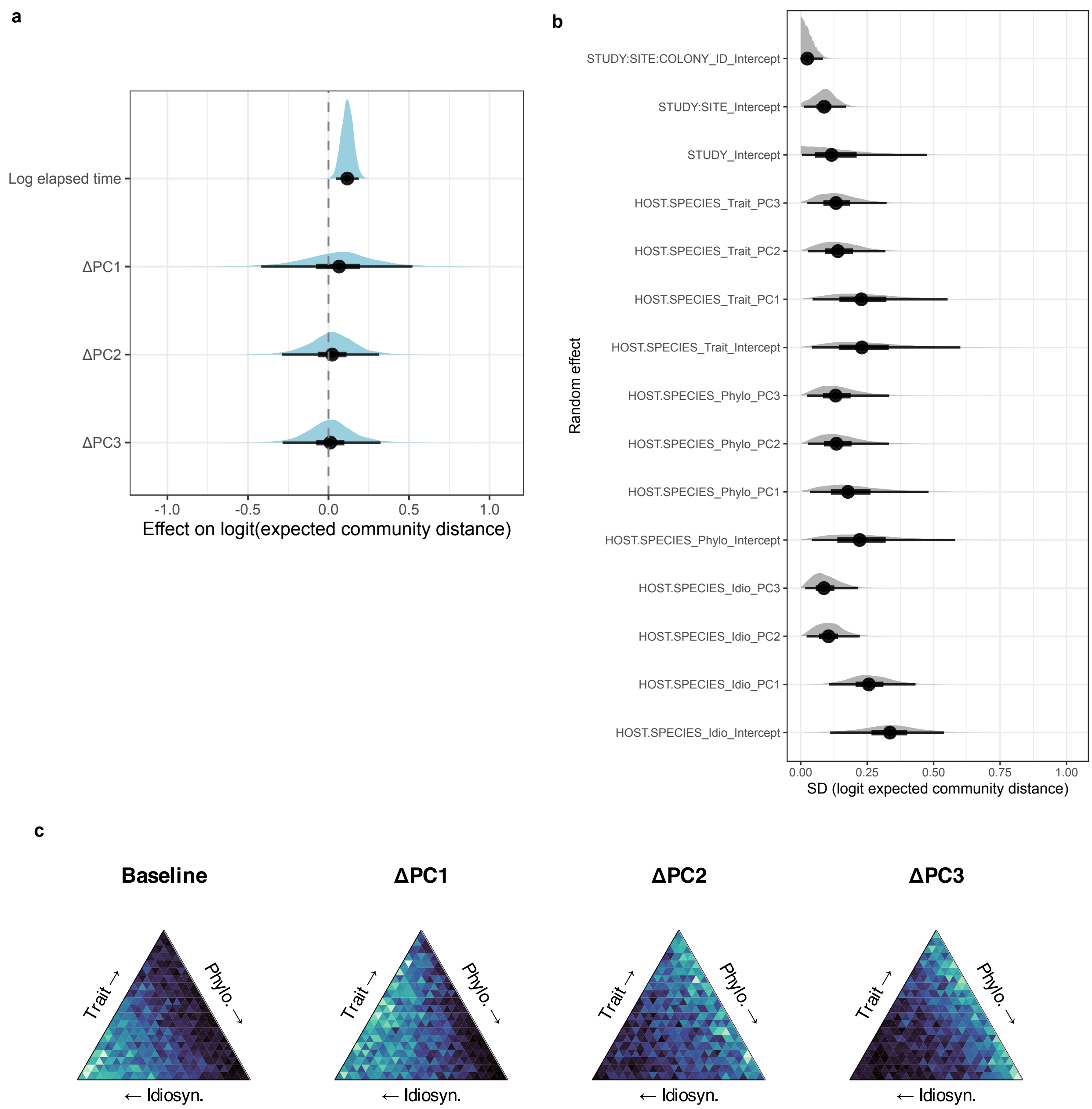


**Figure S2.3** Weighted-UniFrac sensitivity analysis of DIV-level community turnover. (a) Posterior distributions of fixed effects. (b) Posterior distributions of random-effect standard deviations. (c) Posterior allocation of species-level variance among phylogenetic, trait-associated, and idiosyncratic components for baseline turnover and sensitivity to ΔPC1-ΔPC3.

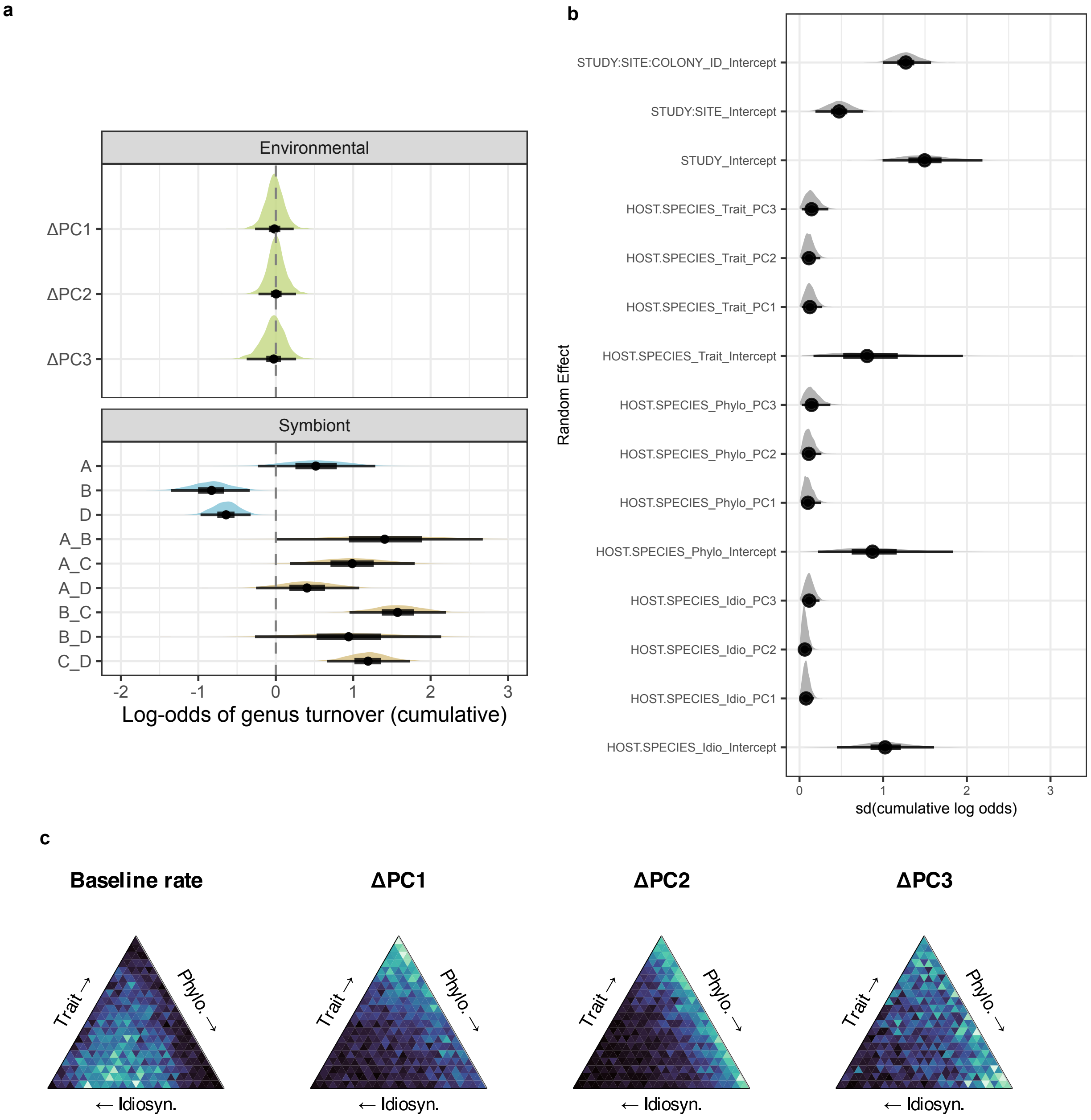


**Figure S2.4** Genus-level ordinal sensitivity analysis excluding colonies first observed in a mixed-genus state. (a) Posterior distributions of fixed effects. (b) Posterior distributions of random-effect standard deviations. (c) Posterior allocation of species-level variance among phylogenetic, trait-associated, and idiosyncratic components for baseline turnover and sensitivity to ΔPC1-ΔPC3.

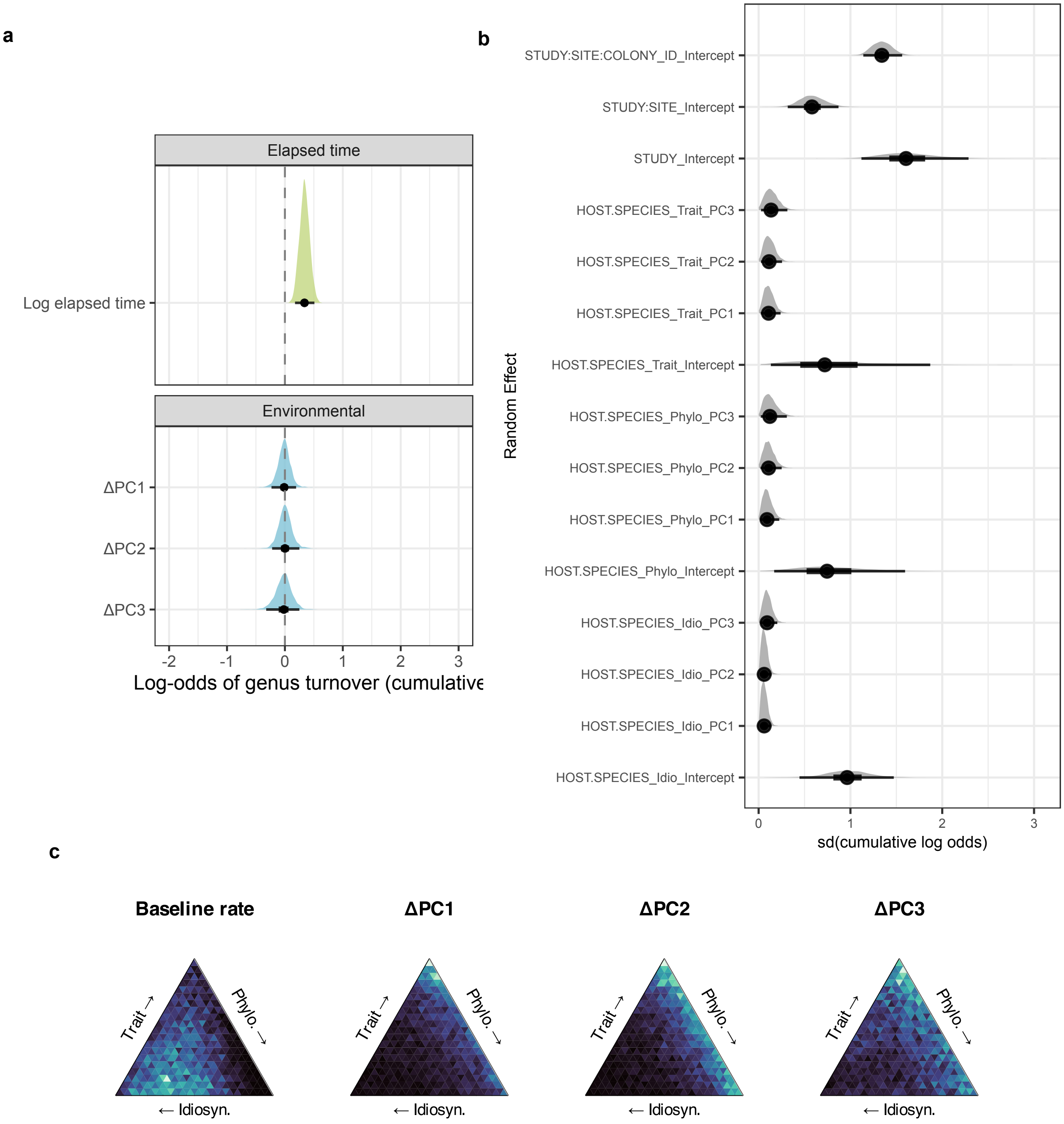


**Figure S2.5** Genus-level ordinal sensitivity analysis omitting prior symbiont genus from the fixed-effect design matrix while retaining all otherwise eligible observations (a) Posterior distributions of fixed effects. (b) Posterior distributions of random-effect standard deviations. (c) Posterior allocation of species-level variance among phylogenetic, trait-associated, and idiosyncratic components for baseline turnover and sensitivity to ΔPC1-ΔPC3.

#### Supplemental Tables

**Table S2.1** Reduced-timeframe genus-level ordinal model. Posterior medians and 95% credible intervals are shown for fixed effects and random-effect standard deviations. Fixed effects are on the cumulative-logit scale.

##### Fixed Effects

| *Term* | *Median* | *2.5%* | *97.5%* |
|---|---|---|---|
| *Log elapsed time* | 0.276 | 0.084 | 0.474 |
| *ΔPC1* | -0.017 | -0.362 | 0.308 |
| *ΔPC2* | -0.008 | -0.336 | 0.307 |
| *ΔPC3* | -0.060 | -0.513 | 0.353 |
| *Prior symbiont state: Symbiodinium* | 2.571 | 1.737 | 3.382 |
| *Prior symbiont state: Symbiodinium + Cladocopium* | 2.519 | 1.817 | 3.212 |
| *Prior symbiont state: Symbiodinium + Cladocopium + Durusdinium* | 0.812 | -0.733 | 2.340 |
| *Prior symbiont state: Symbiodinium + Cladocopium + Gerakladium* | 0.868 | -0.680 | 2.411 |
| *Prior symbiont state: Symbiodinium + Durusdinium* | 0.825 | 0.259 | 1.359 |
| *Prior symbiont state: Breviolum* | -0.819 | -1.704 | 0.023 |
| *Prior symbiont state: Breviolum + Cladocopium* | 2.119 | 1.378 | 2.851 |
| *Prior symbiont state: Breviolum + Cladocopium + Durusdinium* | 0.388 | -1.337 | 2.101 |
| *Prior symbiont state: Breviolum + Durusdinium* | 1.174 | -0.124 | 2.579 |
| *Prior symbiont state: Cladocopium + Durusdinium* | 1.349 | 1.027 | 1.671 |
| *Prior symbiont state: Cladocopium + Gerakladium* | 0.272 | -1.430 | 2.093 |
| *Prior symbiont state: Durusdinium* | -0.785 | -1.128 | -0.460 |
| *Intercept[1]: no genus turnover vs. codominance or major turnover* | 2.040 | 0.976 | 3.008 |
| *Intercept[2]: no turnover or codominance vs. major turnover* | 4.176 | 3.084 | 5.167 |

##### Random Effects

| *Random effect* | *Median* | *2.5%* | *97.5%* |
|---|---|---|---|
| *Species phylogenetic component: intercept* | 0.697 | 0.166 | 1.488 |
| *Species phylogenetic component: ΔPC1 slope* | 0.135 | 0.025 | 0.333 |
| *Species phylogenetic component: ΔPC2 slope* | 0.150 | 0.031 | 0.349 |
| *Species phylogenetic component: ΔPC3 slope* | 0.170 | 0.034 | 0.461 |
| *Species idiosyncratic component: intercept* | 0.767 | 0.321 | 1.282 |
| *Species idiosyncratic component: ΔPC1 slope* | 0.112 | 0.022 | 0.251 |
| *Species idiosyncratic component: ΔPC2 slope* | 0.084 | 0.015 | 0.194 |
| *Species idiosyncratic component: ΔPC3 slope* | 0.151 | 0.027 | 0.350 |
| *Species trait component: intercept* | 0.734 | 0.147 | 1.725 |
| *Species trait component: ΔPC1 slope* | 0.161 | 0.034 | 0.367 |
| *Species trait component: ΔPC2 slope* | 0.168 | 0.036 | 0.366 |
| *Species trait component: ΔPC3 slope* | 0.206 | 0.043 | 0.495 |
| *Study intercept* | 1.388 | 0.865 | 2.170 |
| *Site-within-study intercept* | 0.445 | 0.219 | 0.714 |
| *Colony-within-site intercept* | 0.607 | 0.148 | 0.941 |

**Table S2.2** First-turnover Bernoulli model. Posterior medians and 95% credible intervals are shown for fixed effects and random-effect standard deviations. Fixed effects are on the log-odds scale.

#### Fixed Effects

| *Term* | *Median* | *2.5%* | *97.5%* |
|---|---|---|---|
| *Log elapsed time* | 0.414 | 0.213 | 0.625 |
| *ΔPC1* | -0.037 | -0.310 | 0.214 |
| *ΔPC2* | -0.027 | -0.278 | 0.251 |
| *ΔPC3* | -0.001 | -0.387 | 0.363 |
| *Prior symbiont state: Symbiodinium* | 0.744 | -0.069 | 1.568 |
| *Prior symbiont state: Symbiodinium + Breviolum* | 0.381 | -1.378 | 2.185 |
| *Prior symbiont state: Symbiodinium + Cladocopium* | 2.776 | 1.972 | 3.708 |
| *Prior symbiont state: Symbiodinium + Cladocopium + Durusdinium* | 1.564 | -0.048 | 3.074 |
| *Prior symbiont state: Symbiodinium + Durusdinium* | 0.978 | 0.229 | 1.745 |
| *Prior symbiont state: Breviolum* | -1.132 | -1.807 | -0.503 |
| *Prior symbiont state: Breviolum + Cladocopium* | 3.076 | 2.060 | 4.176 |
| *Prior symbiont state: Breviolum + Cladocopium + Durusdinium* | 0.511 | -1.227 | 2.251 |
| *Prior symbiont state: Breviolum + Durusdinium* | 1.334 | -0.215 | 2.939 |
| *Prior symbiont state: Cladocopium + Durusdinium* | 1.992 | 1.593 | 2.449 |
| *Prior symbiont state: Durusdinium* | -0.655 | -0.995 | -0.327 |
| *Prior symbiont state: Durusdinium + Gerakladium* | 0.386 | -1.448 | 2.277 |

#### Random Effects

| *Random effect* | *Median* | *2.5%* | *97.5%* |
|---|---|---|---|
| *Species phylogenetic component: intercept* | 0.729 | 0.220 | 1.469 |
| *Species phylogenetic component: ΔPC1 slope* | 0.118 | 0.026 | 0.265 |
| *Species phylogenetic component: ΔPC2 slope* | 0.115 | 0.023 | 0.278 |
| *Species phylogenetic component: ΔPC3 slope* | 0.171 | 0.032 | 0.416 |
| *Species idiosyncratic component: intercept* | 0.866 | 0.420 | 1.310 |
| *Species idiosyncratic component: ΔPC1 slope* | 0.069 | 0.014 | 0.162 |
| *Species idiosyncratic component: ΔPC2 slope* | 0.074 | 0.015 | 0.173 |
| *Species idiosyncratic component: ΔPC3 slope* | 0.171 | 0.044 | 0.311 |
| *Species trait component: intercept* | 0.675 | 0.124 | 1.550 |
| *Species trait component: ΔPC1 slope* | 0.130 | 0.027 | 0.274 |
| *Species trait component: ΔPC2 slope* | 0.127 | 0.028 | 0.285 |
| *Species trait component: ΔPC3 slope* | 0.183 | 0.033 | 0.458 |
| *Study intercept* | 1.365 | 0.905 | 1.979 |
| *Site-within-study intercept* | 0.391 | 0.163 | 0.638 |
| *Colony-within-site intercept* | 0.557 | 0.052 | 1.082 |

**Table S2.3** Weighted-UniFrac DIV-level beta model. Posterior medians and 95% credible intervals are shown for fixed effects and random-effect standard deviations. Fixed effects are on the logit scale of expected weighted-UniFrac distance.

##### Fixed Effects

| *Term* | *Median* | *2.5%* | *97.5%* |
|---|---|---|---|
| *Log elapsed time* | 0.117 | 0.046 | 0.187 |
| *ΔPC1* | 0.066 | -0.418 | 0.521 |
| *ΔPC2* | 0.023 | -0.288 | 0.313 |
| *ΔPC3* | 0.012 | -0.285 | 0.323 |
| *Beta mean intercept* | -1.997 | -2.643 | -1.383 |
| *Beta precision (φ)* | 2.956 | 2.763 | 3.154 |

##### Random Effects

| *Random effect* | *Median* | *2.5%* | *97.5%* |
|---|---|---|---|
| *Species phylogenetic component: intercept* | 0.221 | 0.041 | 0.581 |
| *Species phylogenetic component: ΔPC1 slope* | 0.178 | 0.035 | 0.481 |
| *Species phylogenetic component: ΔPC2 slope* | 0.135 | 0.027 | 0.332 |
| *Species phylogenetic component: ΔPC3 slope* | 0.131 | 0.025 | 0.333 |
| *Species idiosyncratic component: intercept* | 0.336 | 0.111 | 0.539 |
| *Species idiosyncratic component: ΔPC1 slope* | 0.256 | 0.107 | 0.432 |
| *Species idiosyncratic component: ΔPC2 slope* | 0.105 | 0.022 | 0.222 |
| *Species idiosyncratic component: ΔPC3 slope* | 0.087 | 0.017 | 0.216 |
| *Species trait component: intercept* | 0.230 | 0.042 | 0.601 |
| *Species trait component: ΔPC1 slope* | 0.228 | 0.044 | 0.553 |
| *Species trait component: ΔPC2 slope* | 0.140 | 0.027 | 0.318 |
| *Species trait component: ΔPC3 slope* | 0.133 | 0.025 | 0.323 |
| *Study intercept* | 0.116 | 0.005 | 0.475 |
| *Site-within-study intercept* | 0.088 | 0.011 | 0.171 |
| *Colony-within-site intercept* | 0.025 | 0.001 | 0.084 |

**Table S2.4** Genus-level ordinal model omitting initially co-dominant colonies. Posterior medians and 95% credible intervals are shown for fixed effects and random-effect standard deviations. Fixed effects are on the cumulative-logit scale.

#### Fixed Effects

| ***Term*** | ***Median*** | ***2.5%*** | ***97.5%*** |
|---|---|---|---|
| *Log elapsed time* | 0.398 | 0.202 | 0.597 |
| *ΔPC1* | -0.017 | -0.267 | 0.232 |
| *ΔPC2* | 0.004 | -0.223 | 0.262 |
| *ΔPC3* | -0.026 | -0.375 | 0.263 |
| *Prior symbiont state: Symbiodinium* | 0.517 | -0.231 | 1.285 |
| *Prior symbiont state: Symbiodinium + Breviolum* | 1.414 | 0.017 | 2.785 |
| *Prior symbiont state: Symbiodinium + Cladocopium* | 0.987 | 0.184 | 1.794 |
| *Prior symbiont state: Symbiodinium + Cladocopium + Durusdinium* | 0.593 | -0.843 | 1.980 |
| *Prior symbiont state: Symbiodinium + Cladocopium + Gerakladium* | 0.957 | -0.704 | 2.586 |
| *Prior symbiont state: Symbiodinium + Durusdinium* | 0.403 | -0.255 | 1.078 |
| *Prior symbiont state: Breviolum* | -0.829 | -1.354 | -0.337 |
| *Prior symbiont state: Breviolum + Cladocopium* | 1.573 | 0.952 | 2.197 |
| *Prior symbiont state: Breviolum + Cladocopium + Durusdinium* | 0.391 | -1.391 | 2.157 |
| *Prior symbiont state: Breviolum + Durusdinium* | 0.941 | -0.268 | 2.138 |
| *Prior symbiont state: Cladocopium + Durusdinium* | 1.192 | 0.660 | 1.734 |
| *Prior symbiont state: Cladocopium + Gerakladium* | 0.530 | -1.285 | 2.255 |
| *Prior symbiont state: Durusdinium* | -0.642 | -0.971 | -0.325 |
| *Intercept[1]: no genus turnover vs. codominance or major turnover* | 2.697 | 1.514 | 3.710 |
| *Intercept[2]: no turnover or codominance vs. major turnover* | 4.011 | 2.836 | 5.040 |

#### Random Effects

| ***Random effect*** | ***Median*** | ***2.5%*** | ***97.5%*** |
|---|---|---|---|
| *Species phylogenetic component: intercept* | 0.873 | 0.222 | 1.834 |
| *Species phylogenetic component: ΔPC1 slope* | 0.101 | 0.020 | 0.255 |
| *Species phylogenetic component: ΔPC2 slope* | 0.111 | 0.025 | 0.261 |
| *Species phylogenetic component: ΔPC3 slope* | 0.144 | 0.026 | 0.369 |
| *Species idiosyncratic component: intercept* | 1.024 | 0.447 | 1.609 |
| *Species idiosyncratic component: ΔPC1 slope* | 0.079 | 0.017 | 0.174 |
| *Species idiosyncratic component: ΔPC2 slope* | 0.063 | 0.013 | 0.150 |
| *Species idiosyncratic component: ΔPC3 slope* | 0.116 | 0.025 | 0.239 |
| *Species trait component: intercept* | 0.808 | 0.167 | 1.953 |
| *Species trait component: ΔPC1 slope* | 0.124 | 0.026 | 0.271 |
| *Species trait component: ΔPC2 slope* | 0.113 | 0.022 | 0.249 |
| *Species trait component: ΔPC3 slope* | 0.144 | 0.027 | 0.345 |
| *Study intercept* | 1.496 | 0.993 | 2.187 |
| *Site-within-study intercept* | 0.473 | 0.190 | 0.761 |
| *Colony-within-site intercept* | 1.271 | 0.995 | 1.572 |

**Table S2.5** Ordinal model without prior genus. Posterior medians and 95% credible intervals are shown for fixed effects and random-effect standard deviations. Fixed effects are on the cumulative-logit scale.

#### Fixed Effects

| *Term* | *Median* | *2.5%* | *97.5%* |
|---|---|---|---|
| *Log elapsed time* | 0.338 | 0.175 | 0.509 |
| *ΔPC1* | -0.012 | -0.231 | 0.193 |
| *ΔPC2* | 0.001 | -0.221 | 0.250 |
| *ΔPC3* | -0.018 | -0.323 | 0.249 |
| *Intercept[1]: no genus turnover vs. codominance or major turnover* | 2.380 | 1.293 | 3.368 |
| *Intercept[2]: no turnover or codominance vs. major turnover* | 4.060 | 2.966 | 5.071 |

#### Random Effects

| *Random effect* | *Median* | *2.5%* | *97.5%* |
|---|---|---|---|
| *Species phylogenetic component: intercept* | 0.746 | 0.169 | 1.595 |
| *Species phylogenetic component: ΔPC1 slope* | 0.092 | 0.017 | 0.224 |
| *Species phylogenetic component: ΔPC2 slope* | 0.108 | 0.020 | 0.251 |
| *Species phylogenetic component: ΔPC3 slope* | 0.122 | 0.023 | 0.307 |
| *Species idiosyncratic component: intercept* | 0.964 | 0.445 | 1.472 |
| *Species idiosyncratic component: ΔPC1 slope* | 0.060 | 0.012 | 0.140 |
| *Species idiosyncratic component: ΔPC2 slope* | 0.061 | 0.012 | 0.142 |
| *Species idiosyncratic component: ΔPC3 slope* | 0.093 | 0.019 | 0.203 |
| *Species trait component: intercept* | 0.721 | 0.133 | 1.869 |
| *Species trait component: ΔPC1 slope* | 0.108 | 0.023 | 0.239 |
| *Species trait component: ΔPC2 slope* | 0.114 | 0.026 | 0.255 |
| *Species trait component: ΔPC3 slope* | 0.135 | 0.026 | 0.313 |
| *Study intercept* | 1.605 | 1.120 | 2.286 |
| *Site-within-study intercept* | 0.581 | 0.317 | 0.870 |
| *Colony-within-site intercept* | 1.342 | 1.141 | 1.563 |
