## Supplemental File 3 for "Within-host algal symbiont turnover is not a general response to environmental change in reef corals"

### Find eligible coral meta-analysis

I want to use the following criteria to do a literature search for papers to include in a met analysis. Do not include any papers already in my meta analysis in the list (attached). The criteria are:

1. The paper must have measured zooxanthellate, scleractianian coral colonies at at least two time points.
2. The paper must have quantified the Symbiodiniaceae community within each colony through time at the genus, species, or strain level.
3. Tagged or genotyped coral colonies must have been tracked, so final measurements are at the colony-level.
4. Colonies must have been measured on the reef and could not have undergone any experimental manipulation, such as reciprocal transplants or common garden treatments. Control groups for these types of experiments which were not manipulated can be included.

For each paper, give me the full citation. Be sure to double check your citations and paper links so I can verify them manually. If the paper lists public data accessions (github/NCBI) list them in the table in their own columns.

Already Included Studies:

STUDY SPECIES COLONIES OBS REGION (SITES) DATES

(THORNHILL, FITT, ET AL., 2006)

6 126 1728 Northern Caribbean, Florida & Bahamas (3) 1998-2004

(THORNHILL, LAJEUNESSE, ET AL., 2006)

3 24 150 Northern Caribbean, Florida & Bahamas (7) 2002-2005

(JONES ET AL., 2008)

1 127 254 Great Barrier Reef (1) 2004-2006

(SAMPAYO ET AL., 2008)

1 7 20 Great Barrier Reef (1) 2005-2006

(STAT ET AL., 2009)

10 139 417 Great Barrier Reef (1) 2001-2002

(THORNHILL ET AL., 2009)

2 50 200 Northern Caribbean, Florida & Bahamas (5) 2003-2005

(KIMES ET AL., 2013)

1 6 12 SE Caribbean (1) 2007

(ROUZÉ ET AL., 2016)

1 11 84 Central South Pacific (2) 2011-2012

(MANZELLO ET AL., 2019)

1 124 248 Northern Caribbean, Florida & Bahamas (3) 2015-2016

(CARBALLO-BOLAÑOS ET AL., 2019)

1 25 75 NE Asia (2) 2016-2017

(EPSTEIN ET AL., 2019)

1 10 88 Great Barrier Reef (1) 2016-2017

(ROUZÉ ET AL., 2019)

3 45 360 Central South Pacific (4) 2011-2012

(CLAAR ET AL., 2020)

3 182 605 Eastern Micronesia (15) 2014-2019

(HOWELLS ET AL., 2020)

3 118 375 NW Arabian Sea (4) 2012-2014

(JANDANG ET AL., 2022)

2 32 81 SE Asia (1) 2018

(LEINBACH ET AL., 2023)

1 26 50 Central South Pacific (1) 2019

(MCRAE ET AL., 2023)

3 30 111 NE Asia (2) 2018-2019

(NÚÑEZ-PONS ET AL., 2023)

3 127 501 Hawaiian Islands (1) 2014-2015

(PALACIO-CASTRO ET AL., 2023)

3 29 87 Eastern Tropical Pacific (1) 2014-2016

(QUEK ET AL., 2023)

3 24 144 SE Asia (2) 2018-2020

(STARKO ET AL., 2023)

1 163 524 Eastern Micronesia (14) 2014-2019

(DE SOUZA ET AL., 2023)

1 468 936 Hawaiian Islands (1) 2018-2019

(CUNNING ET AL., 2024)

9 56 141 SE Caribbean (3) 2017-2019

(LOCK ET AL., 2024)

1 19 30 Western Micronesia (1) 2018

(TERRANEO ET AL., 2024)

4 62 180 SW Pacific (4) 2016

(HUFFMYER ET AL., 2025)

3 125 421 Central South Pacific (3) 2020

(LI ET AL., 2025)

1 5 20 SE Asia (1) 2023-2024

(TURNHAM ET AL., 2025)

13 32 146 Western Micronesia (1) 2013-2022

(VAN NYNATTEN ET AL., 2025)

2 237 605 Eastern Micronesia (16) 2013-2023

(POWELL ET AL., 2026)

3 33 81 SE Caribbean (4) 2020-2021

(GLASS ET AL., 2026)

1 163 440 Mesoamerica (4) 2022-2024

TOTAL 59 2625 9114 14 Regions, 95 Sites 1998-2024

► 15 previous messages

#### Additional eligible papers

I found nine papers not present in your supplied list that satisfy all four criteria. I treated legacy Symbiodinium clades C and D as genus-level identifications because they correspond to the modern genera *Cladocopium* and *Durusdinium*.

| Full citation and verified link | Longitudinal evidence | Symbiodiniaceae resolution | GitHub | NCBI access |
|---|---|---|---|---|

| | | | | |
|---|---|---|---|---|
| Chen, C. A., Wang, J.-T., Fang, L.-S., & Yang, Y.-W. (2005). Fluctuating algal symbiont communities in *Acropora palifera* (Scleractinia: Acroporidae) from Taiwan. *Marine Ecology Progress Series, 295*, 113–121. DOI (https://doi.org/10.3354/meps295113) | Twenty colonies tagged; eight surviving tagged colonies sampled bimonthly, March 2000–July 2001, on the reef. | PCR-RFLP: legacy clades C and D | None stated | None stated |
| Chen, C.-P., Tseng, C.-H., Chen, C. A., & Tang, S.-L. (2011). The dynamics of microbial partnerships in the coral *Isopora palifera*. *The ISME Journal, 5*(4), 728–740. DOI (https://doi.org/10.1038/ismej.2010.151); PMC full text (https://pmc.ncbi.nlm.nih.gov/articles/PMC3105734/) | Three tagged colonies sampled during seven months of 2008 under natural reef conditions. | qPCR: legacy clades C and D | None stated | None stated for coral-symbiont d[illegible] |
| LaJeunesse, T. C., Smith, R., Walther, M., Pinzón, J., Pettay, D. T., McGinley, M., Aschaffenburg, M., Medina-Rosas, P., Cupul-Magaña, A. L., Pérez, A. L., Reyes-Bonilla, H., & Warner, M. E. (2010). Host–symbiont recombination versus natural selection in the response of coral–dinoflagellate symbioses to environmental disturbance. *Proceedings of the Royal Society B: Biological Sciences, 277*, 2925–2934. DOI (https://doi.org/10.1098/rspb.2010.0385); PMC full text (https://pmc.ncbi.nlm.nih.gov/articles/PMC2982020/) | Eighty-four tagged *Pocillopora* colonies along permanent transects; repeated sampling around a natural cold-water bleaching event. | ITS2-DGGE/sequence types C1b-c and D1 | None stated | None stated |
| Pettay, D. T., Wham, D. C., Pinzón, J. H., & LaJeunesse, T. C. (2011). Genotypic diversity and spatial–temporal distribution of *Symbiodinium* clones in an abundant reef coral. *Molecular Ecology, 20*(24), 5197–5212. DOI (https://doi.org/10.1111/j.1365-294X.2011.05357.x); PMC full text (https://pmc.ncbi.nlm.nih.gov/articles/PMC5957298/) | Nine distinctive *Pocillopora* colonies relocated and resampled after nine months; matching within-colony orientations sampled. | Multilocus microsatellite genotypes of *Durusdinium glynnii* strains | None stated | None stated |
| Hsu, C.-M., Keshavmurthy, S., Denis, V., Kuo, C.-Y., Wang, J.-T., Meng, P.-J., & Chen, C. A. (2012). Temporal and spatial variations in symbiont communities of catch bowl coral *Isopora palifera* (Scleractinia: Acroporidae) on reefs in Kenting National Park, Taiwan. *Zoological Studies, 51*(8), 1343–1353. Verified full-text PDF (https://researchonline.jcu.edu.au/32980/1/32980%20Hsu%20et%20al%202012.pdf) | Thirty-two tagged colonies sampled bimonthly, March 2006–November 2009. | 28S RFLP and ITS2-DGGE: C3 and D1a | None stated | None stated |
| Kemp, D. W., Hernandez-Pech, X., Iglesias-Prieto, R., Fitt, W. K., & Schmidt, G. W. (2014). Community dynamics and physiology of *Symbiodinium* spp. before, during, and after a coral bleaching event. *Limnology and Oceanography, 59*(3), 788–797. DOI (https://doi.org/10.4319/lo.2014.59.3.0788) | Tagged *Orbicella faveolata* followed before, during and after natural bleaching; pigmented and bleached regions of the same colonies | ITS2 types A3, B17, C7 and D1a | None stated | None stated |

| | | | | |
|---|---|---|---|---|
| | were compared. | | | |
| Quigley, K. M., Willis, B. L., & Kenkel, C. D. (2019). Transgenerational inheritance of shuffled symbiont communities in the coral *Montipora digitata*. *Scientific Reports, 9*, 13328. DOI (https://doi.org/10.1038/s41598-019-50045-y); PMC full text (https://pmc.ncbi.nlm.nih.gov/articles/PMC6746730/) | The same four tagged maternal colonies sampled in 2015 and 2016 across natural bleaching. | ITS2 amplicon sequence variants, including genus- and intragenus-level changes | None stated | SRP077416 (https://www.ncbi.nlm[illegible] term=SRP077416); SRP133660 (https://www.ncbi.nlm.nih.gov/sra[illegible] |
| Thomas, L., López, E. H., Morikawa, M. K., & Palumbi, S. R. (2019). Transcriptomic resilience, symbiont shuffling, and vulnerability to recurrent bleaching in reef-building corals. *Molecular Ecology, 28*(14), 3371–3382. DOI (https://doi.org/10.1111/mec.15143) | The same tracked *Acropora gemmifera* and *A. hyacinthus* colonies sampled at four recovery time points following natural bleaching; survival tracked through a second event. | RNA-seq reads mapped to cp23S, ITS1 and ITS2 symbiont haplotypes | None stated | PRJNA522650 (https://www.ncbi.nlm.nih.gov/bio[illegible] |
| Jung, E. M. U., Stat, M., Thomas, L., Koziol, A., & Schoepf, V. (2021). Coral host physiology and symbiont dynamics associated with differential recovery from mass bleaching in an extreme, macro-tidal reef environment in northwest Australia. *Coral Reefs, 40*(3), 893–905. DOI (https://doi.org/10.1007/s00338-021-02094-x) | Healthy and naturally bleached *Acropora aspera* colonies tagged and resampled in April and November 2016. | Symbiodiniaceae community composition, dominated by *Cladocopium* | None stated | None stated |

### Eligible design, but likely overlaps an included study

Lewis et al. (2024) clearly satisfies the biological criteria, but its Palau colonies and 2013–2022 sampling period appear likely to overlap the already-included Turnham et al. (2025) dataset. I would not add it as independent data without matching colony and sample identifiers:

Lewis, A. M., Butler, C. C., Turnham, K. E., Wham, D. F., Hoadley, K. D., Smith, R. T., Kemp, D. W., Warner, M. E., & LaJeunesse, T. C. (2024). The diversity, distribution, and temporal stability of coral 'zooxanthellae' on a Pacific reef: from the scale of individual colonies to across the host community. *Coral Reefs, 43*(4), 841–856. DOI (https://doi.org/10.1007/s00338-024-02503-x).

Important exclusions included population-level resampling without colony identity (LaJeunesse et al. 2009; Boulotte et al. 2016; Lee et al. 2016) and aquarium, transplant or induced-bleaching experiments. "None stated" means no repository was reported in the article, not that unpublished data cannot exist.

> Can you explain why these weren't included? A reviewer specifically recommended that they should be. Rerun the search after evaluating these papers. You can modify search criteria slightly, but do not return studies that did not track colonies longitudinally, experimentally manipulated coral, or did not track colonies under field conditions.
>
> *Chen C, Wang A, Fang L, Yang Y (2005) Fluctuating algal symbiont communities in Acropora palifera (Scleractinia: Acroporidae) from Taiwan. Marine Ecology-Progress Series 295:113–121*
> *Kemp DW, Hernández-Pech X, Iglesias-Prieto R, Fitt WK, Schmidt GW (2014) Community dynamics and physiology of Symbiodinium spp. before, during, and after a coral beaching event. Limnology and oceanography 59:788–797*
> *Cunning R, Ritson-Williams R, Gates RD (2016) Patterns of bleaching and recovery of Montipora capitata in Kāne'ohe Bay, Hawai'i, USA. Marine Ecology-Progress Series 551:131–139*
> *LaJeunesse TC, Smith R, Walther M, Pinzon J, Pettay DT, McGinley M, Aschaffenburg M, Medina-Rosas P, Cupul-Magana AL, Perez AL, Reyes-Bonilla H, Warner ME (2010) Host-symbiont recombination versus natural selection in the response of coral-dinoflagellate symbioses to environmental disturbance. Proceedings Of The Royal Society Of London Series B-Biological Sciences 277:2925–2934*

► 9 previous messages

The reviewer's four papers all meet the biological criteria and should be included. Three—Chen (2005), Kemp (2014), and LaJeunesse (2010)—were present in my earlier candidate results but not in your existing meta-analysis. Cunning (2016) was a genuine search omission.

I broadened the search in two defensible ways:

- Historical Symbiodinium clades/phylotypes count as genus-level resolution because clades A–D broadly correspond to the modern genera *Symbiodinium*, *Breviolum*, *Cladocopium*, and *Durusdinium*.
- Removing small tissue fragments for observational molecular sampling is not considered experimental manipulation. Where a paper also contains an experiment, only the unmanipulated reef time series is eligible.

### Reviewer-recommended papers

| Paper | Why it qualifies | GitHub | NCBI/GenBank | Other public data |
|---|---|---|---|---|
| Chen, C. A., Wang, J.-T., Fang, L.-S., & Yang, Y.-W. (2005). Fluctuating algal symbiont communities in *Acropora palifera* (Scleractinia: Acroporidae) from Taiwan. *Marine Ecology Progress Series, 295*, 113–121. https://doi.org/10.3354/meps295113 (https://doi.org/10.3354/meps295113) | Directly tracked tagged reef colonies bimonthly from January 2000–July 2001. RFLP phylotyping detected C, D, and mixed C/D communities. No manipulation. | None stated | None stated | None stated |
| LaJeunesse, T. C., Smith, R., Walther, M., Pinzón, J., Pettay, D. T., McGinley, M., Aschaffenburg, M., Medina-Rosas, P., Cupul-Magaña, A. L., Pérez, A. L., Reyes-Bonilla, H., & Warner, M. E. (2010). Host–symbiont recombination versus natural selection in the response of coral–dinoflagellate symbioses to environmental disturbance. *Proceedings of the Royal Society B: Biological Sciences, 277*, 2925–2934. https://doi.org/10.1098/rspb.2010.0385 (https://doi.org/10.1098/rspb.2010.0385) | Tagged *Pocillopora* colonies were resampled following natural cold-water bleaching. ITS2-DGGE resolved colony-level symbiont types through time. | None stated | No consolidated project accession stated | None stated |
| Kemp, D. W., Hernández-Pech, X., Iglesias-Prieto, R., Fitt, W. K., & Schmidt, G. W. (2014). Community dynamics and physiology of *Symbiodinium* spp. before, during, and after a coral bleaching event. *Limnology and Oceanography, 59*(3), 788–797. https://doi.org/10.4319/lo.2014.59.3.0788 (https://doi.org/10.4319/lo.2014.59.3.0788) | Tagged *Orbicella faveolata* colonies were followed before, during, and after natural bleaching in Puerto Morelos. The same colony patches were resampled and ITS2 types—including A3, B17, and C7—were | None stated | None stated | None stated |

| | | | | |
|---|---|---|---|---|
| | measured. | | | |
| Cunning, R., Ritson-Williams, R., & Gates, R. D. (2016). Patterns of bleaching and recovery of *Montipora capitata* in Kāneʻohe Bay, Hawaiʻi, USA. *Marine Ecology Progress Series, 551*, 131–139. https://doi.org/10.3354/meps11733 (https://doi.org/10.3354/meps11733) | Sixty individually tagged reef colonies—naturally bleached and non-bleached—were sampled on six dates from October 2014–April 2015. qPCR quantified clades C and D; ITS2 identified C31 in a subset. No treatment was imposed. | None stated | None stated | Data and scripts (https://doi.org/10.5281/zenodo.53231); environmental records 53226 (https://doi.org/10.5281/zenodo.53226) and 53227 (https://doi.org/10.5281/zenodo.53227) |

Cunning was probably missed because its title emphasizes bleaching and recovery rather than longitudinal symbiont-community monitoring.

### Additional eligible papers found in the rerun

| Full citation | Eligible longitudinal component | GitHub | NCBI/GenBank | Other public data |
|---|---|---|---|---|
| Suwa, R., Hirose, M., & Hidaka, M. (2008). Seasonal fluctuation in zooxanthellar genotype composition and photophysiology in the corals *Pavona divaricata* and *P. decussata*. *Marine Ecology Progress Series, 361*, 129–137. https://doi.org/10.3354/meps07372 (https://doi.org/10.3354/meps07372) | Repeated field measurements of identifiable colonies over a year resolved clades C, D, and mixed C/D communities. Include only the unmanipulated reef time series; exclude the separate laboratory temperature/light exposures. | None stated | None stated | None stated |
| Hume, B. C. C., D'Angelo, C., Smith, E. G., Stevens, J. R., Burt, J., & Wiedenmann, J. (2015). *Symbiodinium thermophilum* sp. nov., a thermotolerant symbiotic alga prevalent in corals of the world's hottest sea, the Persian/Arabian Gulf. *Scientific Reports, 5*, 8562. https://doi.org/10.1038/srep08562 (https://doi.org/10.1038/srep08562) | Three tagged colonies of each of six scleractinian species were sampled at eight field time points over 22 months. ITS2-DGGE resolved A1, C1, C3, C15, D1–4 and other types. Use the Saadiyat reef series; exclude separate aquarium-derived samples used for supplementary phylogenetic work. | None stated | KM487747–KM487749; KP234522–KP234524; KM458273–KM458294 | Dryad (https://doi.org/10.5061/dryad.q75p2) |

### Publications requiring dataset deduplication

These satisfy the design but should not automatically become independent meta-analytic studies:

- Lewis, A. M., Butler, C. C., Turnham, K. E., Hoadley, K. D., Smith, R. T., Kemp, D. W., Wham, D. F., Warner, M. E., & LaJeunesse, T. C. (2024). The diversity, distribution, and temporal stability of coral 'zooxanthellae' on a Pacific reef: from the scale of individual colonies to across the host community. *Coral Reefs, 43*, 841–856. https://doi.org/10.1007/s00338-024-02503-x (https://doi.org/10.1007/s00338-024-02503-x). Its Palau 2013–2022 colony cohort appears to overlap your included Turnham et al. (2025) study.
- Buzzoni, D., Van Nynatten, A., Cunning, R., & Baum, J. K. (2026). Persistent legacy effects of marine heatwaves on coral symbioses. *Global Change Biology, 32*(3), e70818. https://doi.org/10.1111/gcb.70818 (https://doi.org/10.1111/gcb.70818). Its 237 Kiritimati colonies and 2014–2023 sampling almost certainly overlap Van Nynatten et al. (2025). Data: NCBI PRJNA1294313 (https://www.ncbi.nlm.nih.gov/bioproject/PRJNA1294313), GitHub (https://github.com/baumlab/Buzzoni_et_al_2026_Platygyra), and Zenodo (https://doi.org/10.5281/zenodo.18792037).

Therefore, the corrected search adds at least Cunning (2016), Suwa et al. (2008), and Hume et al. (2015), while confirming that Chen (2005), LaJeunesse (2010), and Kemp (2014) are eligible additions to your attached meta-analysis.

> Can you see if the individual level colony data was reported from this paper in any subsequent papers?
>
> # Seasonal fluctuation in zooxanthellar genotype composition and photophysiology in the corals *Pavona divaricata* and *P. decussata*
>
> It’s only aggregated to the opulation level within.

► 6 previous messages

I could not find any subsequent paper or public repository that reports the Suwa et al. (2008) colony-by-date data.

I checked its 51 citing publications, author-linked records, and searches across Dryad, Zenodo, Figshare, and institutional repositories.

The only closely related record is:

Suwa, R., Hirose, M., & Hidaka, M. (2007). *Divergent fluctuations in zooxanthella genotypes and physiological characteristics in the two species of genus Pavona*. University of the Ryukyus research report. Repository record (https://u-ryukyu.repo.nii.ac.jp/records/2001060) · PDF (https://u-ryukyu.repo.nii.ac.jp/record/2001060/files/2006PG-4.pdf)

However:

- It predates the journal paper and appears to be a preliminary presentation of the same study.
- The repository supplies only the report PDF, not a spreadsheet or colony-level dataset.
- I found no evidence that it provides the complete colony × sampling-date matrix.
- Later citing papers—including Lee et al. (2016), Rouzé et al. (2019), and Lewis et al. (2024)—cite the conclusions but analyze independent colonies and do not reproduce Suwa’s observations.

The recoverable information is limited to statements such as three *P. divaricata* colonies remaining clade D and seven showing some C/D change. That is insufficient to reconstruct individual longitudinal trajectories.

Therefore, I would mark Suwa et al. (2008) as:

**Biologically eligible, but unusable for the meta-analysis because colony-level longitudinal outcomes are unavailable.**

The remaining route would be a direct data request to Ryota Suwa, Mamiko Hirose, or Michio Hidaka.

> Did your search look for keywords like “symbiont dynamics”? If not, rerun to check for additional papers. If so, there is no need for another search.

► 5 previous messages

No—the exact phrase “symbiont dynamics” was not previously a primary search term. The targeted rerun found three additional eligible papers absent from your included-study table:

| Study | Eligibility | Public data |
|---|---|---|
| Alessi C, Lemonnier H, Camp EF, Wabete N, Payri C, Rodolfo-Metalpa R (2024). *Algal symbiont diversity in Acropora muricata from the extreme reef of Bouraké associated with resistance to coral bleaching.* **PLOS ONE 19:** e0296902. Article/DOI (https://doi.org/10.1371/journal.pone.0296902) | Tagged natural colonies sampled during bleaching and four months later; ITS2/SymPortal profiles. A separate nubbin was used in a growth experiment, but the longitudinal symbiont observations came from the untransplanted source colonies. | Dryad (https://doi.org/10.5061/dryad.05qfttf8z); NCBI PRJNA1020910 (https://www.ncbi.nlm.nih.gov/bioproject/PRJNA1020910) |
| Lee LK, Leaw CP, Gu H, Lim PT, Tan CH (2026). *Shift of algal symbiont communities in the coral Acropora digitifera: Shuffling and switching during thermal stress and recovery.* **Regional Studies in Marine Science 101:** 105266. Article/DOI (https://doi.org/10.1016/j.rsma.2026.105266) | Tagged reef colonies repeatedly sampled for seven months following natural thermal stress; ITS2 amplicon sequencing; no manipulation reported. | NCBI PRJNA1516876 (https://www.ncbi.nlm.nih.gov/bioproject/PRJNA1516876), SRA study SRP729419 |
| Asadi Gharabaghi S, Collard O, Flot J-F, Ben Hamadou R, Beji M, Shokri MR (2026). *Living on the edge: Host and habitat-associated Symbiodiniaceae dynamics in corals of the world’s hottest sea (the Persian Gulf).* **Marine Environmental Research 222:** 108464. Article/DOI (https://doi.org/10.1016/j.marenvres.2026.108464) | Seasonal ITS2 measurements from the same tagged field colonies; three colonies per species; no experimental treatment reported. Published online 14 October 2026. | No GitHub or NCBI accession located in the currently indexed record. |

The rerun also recovered Jung et al. (2021), but that paper had already appeared in my earlier candidate results:

Jung et al. 2021 (https://doi.org/10.1007/s00338-021-02094-x), with data at OSF (https://doi.org/10.17605/OSF.IO/ZGUDR).

Other prominent hits were either already in your meta-analysis or failed colony-level longitudinal screening.
